# Attractor dynamics of working memory explain a concurrent evolution of stimulus-specific and decision-consistent biases in visual estimation

**DOI:** 10.1101/2023.06.28.546818

**Authors:** Hyunwoo Gu, Joonwon Lee, Sungje Kim, Jaeseob Lim, Hyang-Jung Lee, Heeseung Lee, Min Jin Choe, Dong-gyu Yoo, Jun Hwan (Joshua) Ryu, Sukbin Lim, Sang-Hun Lee

## Abstract

Sensory evidence tends to be fleeting, often unavailable when we categorize or estimate world features. To overcome this, our brains sustain sensory information in working memory. Although keeping that information accurate while acting on it is vital, humans display two canonical biases: estimates are biased toward a few stimuli (“stimulus-specific bias”) and prior decisions (“decision-consistent bias”). Integrative—especially neural mechanistic—accounts of these biases remain scarce. Here, we identify drift dynamics toward discrete attractors as a common source of both biases in orientation estimation, with decisions further steering memory states. Behavior and neuroimaging data reveal how these biases co-evolve through the decision-steered attractor dynamics. Task-optimized recurrent neural networks suggest neural mechanisms that enable categorical decisions to emerge from working memory for continuous stimuli while updating its trajectory, warping decision-consistent biases under stimulus-specific drift.

## Introduction

Adapting to our surroundings engages us in various perceptual tasks on the same object^1,2^, like judging its categorical state (e.g., whether an apple is ‘small’ or ‘large’) and estimating its exact state (e.g., ‘precise size’ of an apple). These tasks often occur in succession, with cognitive processes for earlier tasks influencing later ones. *Decision-consistent bias* is a prime example, where our estimate of a feature aligns with the categorical state of our previous decision (e.g., after deciding on ‘large’, size estimates tend to be ‘larger’ than the actual size). Understanding this can provide insights into the brain’s flexible use of feature representations under varying task demands. While research has clarified how categorical decisions are formed from sensory evidence^3–7^, how the decision-forming process influences the subsequent retention of sensory evidence for future reuse remains unclear.

Decision-consistent bias, once viewed as a perceptual illusion caused by biased readouts of unbiased sensory representations^2^, has been recently reconceptualized^8–11^ as involving post-perceptual processes. In these studies^2,8–11^, since sensory inputs from target stimuli are no longer available during a subsequent estimation task, sensory evidence must be held in working memory (WM)^12^. However, efforts to explain decision-consistent bias in the context of WM have been surprisingly scarce, especially given WM’s dynamic nature^13–15^ and its close relationship with decision-making (DM)^15–18^.

Alongside decision-consistent bias, perceptual estimation exhibits another prominent bias: estimates tend to cluster around specific points in a feature space (e.g., position estimates deviate from cardinal to oblique meridians). This phenomenon, called *stimulus-specific bias*, is common across various domains, including spatial position^19,20^, motion direction^21,22^, orientation^23,24^, and color^25,26^. Although Bayesian approaches have yielded normative accounts of why stimulus-specific bias occurs^27–29^, our understanding of its neural-mechanistic origins remains limited. Moreover, its relationship with decision-consistent bias has not been explored, despite both occurring in similar estimation tasks. These gaps underscore the need for an integrated, mechanistic-level account to clarify their coexistence and interaction in a dynamic context. Focusing on only one bias might ignore the interconnected roots linked to the other.

We hypothesized two intrinsic dynamics of WM that critically influence the co-evolution and interaction of the two biases. Memory states may gradually and randomly shift in a feature space, engendering diffusive representations that grow noisier yet remain unbiased^13,14,30^ (**Figure 1A**). Alternatively, memory states may not only diffuse but also drift toward a few stable points (attractors), engendering drifting and diffusive representations that become increasingly biased and noisy^26,31–33^ (**Figure 1B**). Assuming decision-formation processes steer ongoing memory states in the choice-consistent direction (**Figures 1C,D**, arrows), we expect that the two biases will undergo different time courses depending on whether WM dynamics are driven by diffusion alone (**Figure 1C**) or by both diffusion and drift (**Figure 1D**), and when decisions are made (**Figures 1C,D**, top versus bottom panels).

**Figure 1.**
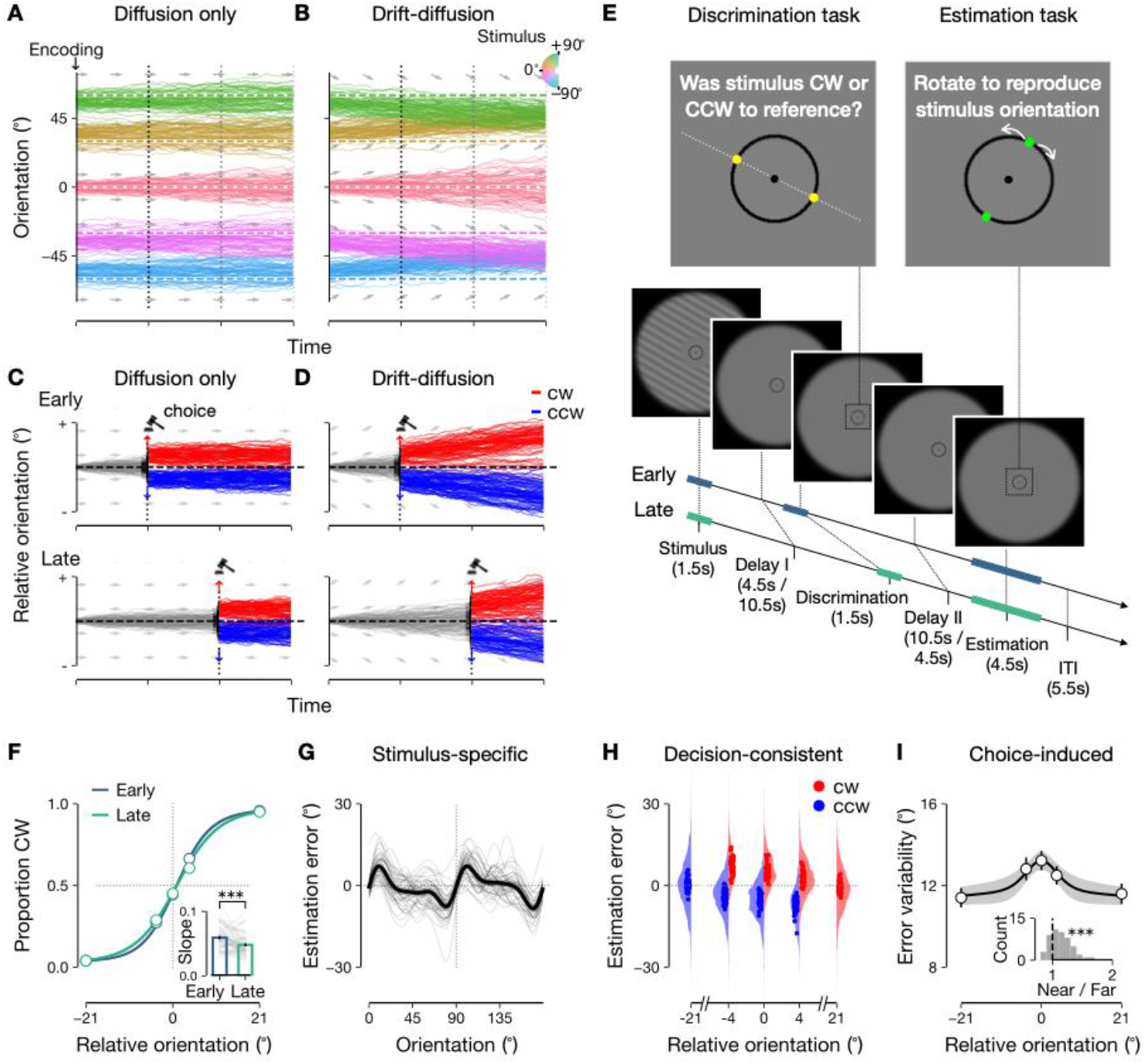
Probing the impact of WM dynamics on behavioral biases. **A-D**, Single-trial memory trajectories simulated under diffusion-only (**A,C**) and drift-diffusion (**B,D**) dynamics. Gray arrows indicate drift directions. **A,B**, Impact on stimulus-specific bias. **C,D**, Impact on decision-consistent bias. Decision timing (gravel) varies between top (early) and bottom (late) panels. **E**, Task paradigm. Top, Yellow and green dots serve as the reference for discrimination and the response frame for estimation; dotted lines, arrows, and white text are shown for illustration purposes. Bottom, Blue and teal bars represent the task epochs in early and late DM trials. **F**, Psychometric curves of discrimination, pooled across individuals. Late DM shows shallower slopes (inset: thin lines, individuals; error bars, s.e.m.; paired t-test, p = 0.0008). **G-I**, Behavioral signatures of stimulus-specific (**G**), decision-consistent (**H**), and choice-induced (**I**) biases. **G**, Estimation errors captured via von Mises function derivatives: thin, individuals; thick, mean across participants ± s.e.m.. **H**, Choice-conditioned distributions of estimation errors: dots, means for individuals; patches, pooled densities. **i**, Error variability measured as inter-quartile range: circles with error bars, mean ± s.e.m.; curve with a gray shadow, Gaussian fit to the data with s.e.m.; inset, variability ratio histogram, with a significant tendency above 1, Wilcoxon signed-rank test, p < 10^−5^. ***, p < 0.001.

To examine these expectations, we designed a task in which participants made sequential categorical choices and point estimates about a remembered stimulus orientation, with varying intervening delays (**Figure 1E**). Monitoring human individuals’ task performance through their choices and estimates while decoding orientation memory from functional magnetic resonance imaging (fMRI) of their visual cortex, we tracked the behavioral and neural signatures of both biases over a prolonged delay period. To further examine whether simple neural mechanisms could recapitulate WM and DM interaction in humans, we also trained recurrent neural networks (RNNs) on the same task and analyzed their dynamics.

Our results across behavioral, fMRI, and RNN analyses convergingly indicate ‘decision-steered attractor dynamics of WM’ as the core mechanism underlying the co-evolution of stimulus-specific and decision-consistent biases. Memory states drift toward attractors, influencing categorical decisions, which in turn bias memory trajectories, creating cascading effects that amplify both biases. RNN simulations mirrored human behavior and fMRI dynamics, showing that decisions emerge through modular interactions among three populations: a WM population maintaining orientation memory, and two DM populations competing for choices, with feedback from DM populations steering WM attractor dynamics. Our work provides an integrated neural-mechanistic explanation of the fundamental biases in perceptual estimation and their evolving interaction, which have not been previously addressed.

## Results

### Behavioral signatures of stimulus-specific and decision-consistent biases

During the fMRI scan, participants memorized the orientation of a briefly shown grating and reported it after a 16.5-s delay. To probe the impact of DM on WM, they first performed a discrimination task during the delay (**Figure 1E**, top). In this task, a dot pair around the fixation (‘reference’) appeared, and participants decided, under moderate time pressure (1.5 s), whether the remembered orientation was tilted clockwise (CW) or counterclockwise (CCW) relative to the reference, whose angle was randomly determined. The timing of the discrimination task varied, occurring either 4.5 s or 10.5 s after stimulus offset (**Figure 1E**, bottom). As anticipated from the temporal deterioration of WM^15,18^, discrimination performance declined when tested later, as indicated by a shallower psychometric curve (**Figure 1F**). In the estimation task, participants rotated another dot pair (‘report frame’) to match the remembered orientation, starting from a randomly chosen angle within 180 degrees.

We confirmed stimulus-specific bias in the estimation task: estimates were repelled from cardinal orientations and attracted toward oblique orientations (**Figure 1G**). This well-known phenomenon, called *cardinal repulsion*^23,27,34,35^, showed modest variation in shape and size among individuals.

Decision-consistent bias was also evident when estimation errors were conditioned on discrimination choices, deviating from the reference in line with the choice. This phenomenon, also known as *reference repulsion*^2,9^, was pronounced in trials where the reference orientation was close to the stimulus (−4°, 0°, 4° on the x-axis in **Figure 1H)**. As noted previously^9^, if choices induce a bias in estimates, the marginal error distribution must widen as the stimulus and reference become more similar in orientation (see **Figures S1D-K** for the rationale).

Consistent with this, error variability was greater in near-reference (relative orientation 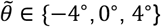) trials compared to far-reference 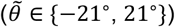 trials (**Figure 1I**). This increased variability was confirmed across various datasets when the reference was relevant to decisions, but not when it served as a distractor (**Figures S1A-C**). We will call this bias *choice-induced bias*, to distinguish it from decision-consistent bias. The latter refers to any deviation aligned with a choice^2,9,10^, measurable from any joint observations of discrete choices and continuous estimates (**Figure 1H**). Choice-induced bias is a particular kind of decision-consistent bias, where commitment to a categorical choice influences estimates beyond what statistical conditioning would predict^8^ (**Figure 1I**).

### Predicting how WM dynamics affect the time courses of biases

We developed phenomenological models to predict how WM dynamics influence bias time courses, with minimal assumptions about sensory encoding and decision commitment’s impact on memory. One model incorporates only diffusion dynamics, allowing random shifts of memory states across trials without bias (**Figures 2A-D**), while the other includes additional drift dynamics that systematically drive memory states in specific directions (**Figures 2E-H**).

**Figure 2.**
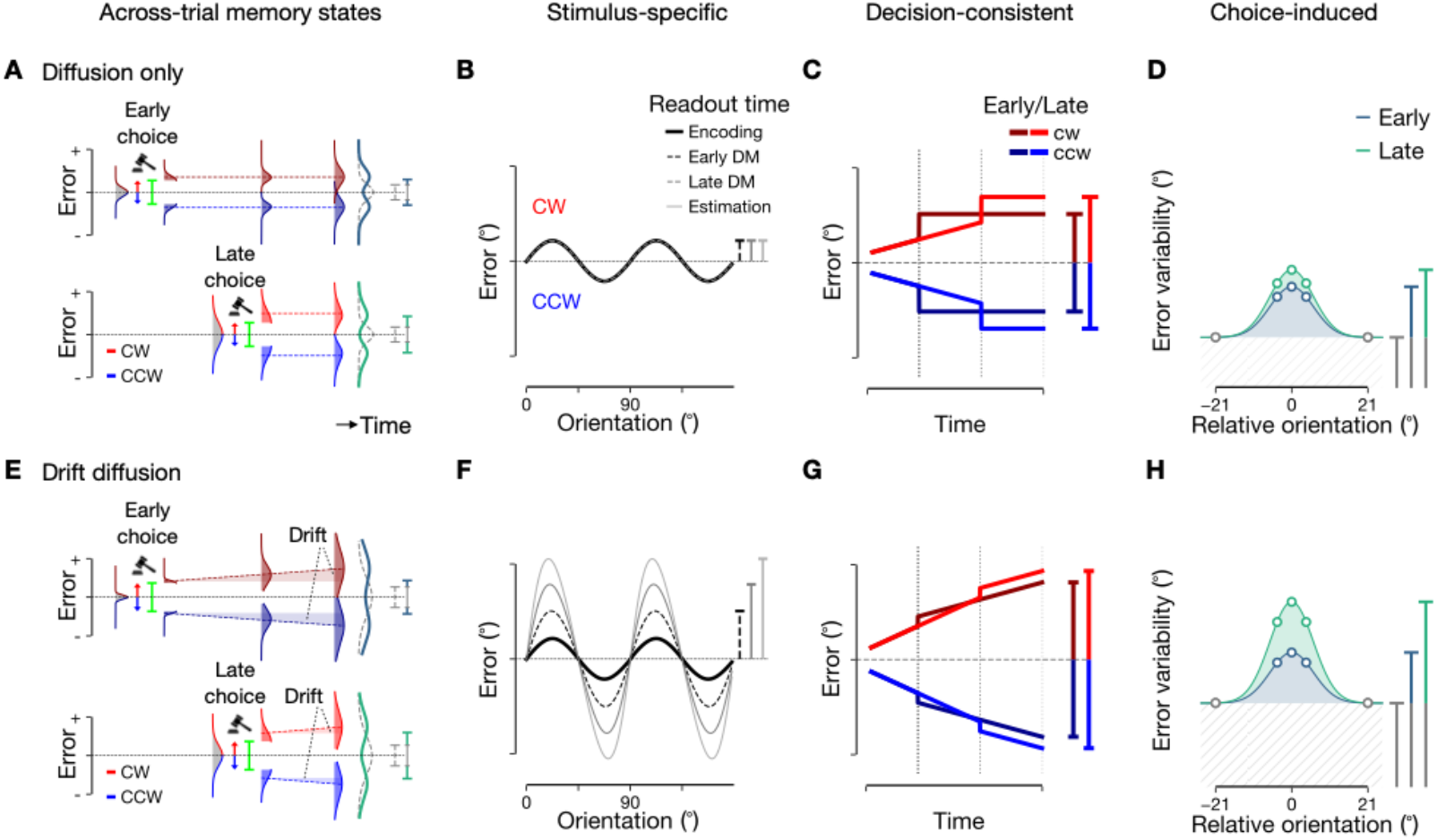
Model predictions of biases under WM dynamics. **A-D**, Diffusion-only model. **E-H**, Drift-diffusion model. **A,E**, Choice-conditioned memory distributions for early (top) and late (bottom) DM trials, with conditional means indicated by horizontal dashed lines. Colored arrows and lime markers represent choice-induced bias during discrimination. Green densities represent marginal error distributions during estimation, alongside dashed densities representing those without choice-induced bias. **B,F**, Stimulus-specific biases across different task epochs. **C,G**, Decision-consistent biases for early and late DM trials. **D,H**, Choice-induced bias captured by near-reference variability for early and late DM trials, alongside dashed gray regions representing error variability without choice-induced bias.

Both models assumed initial orientation memory states follow the efficient encoding principle^27–29^, which allocates more resources to frequently encountered stimuli^36^, leading to higher encoding precision around cardinal orientations. This results in specific error patterns: a mean repulsion away from cardinal orientations and increased variance around oblique ones, aligning with behavioral data. These biases are embedded in the sensory input to the WM system, so memory states already show stimulus-specific bias from the start (**Figures 1A,B**; **2b,f**, darkest curves). To incorporate choice-induced bias into dynamic WM, we also assumed that memory states are abruptly shifted in the chosen direction by a constant amount during the discrimination epoch (vertical arrows in **Figures 1C,D**). This pulse-like shift expands the marginal error distribution for near-reference trials, pushing choice-conditioned distributions apart (**Figures 2A,E**).

Based on these assumptions, we predicted how WM dynamics influence biases in both the diffusion-only and drift-diffusion models. If diffusion solely governs the dynamics (**Figures 1A,2A**), stimulus-specific bias remains unchanged (**Figure 2B**) because stochastic fluctuations of memory states with a zero mean do not cause any systematic deviations (gray arrows in **Figure 1A**). Conversely, decision-consistent bias varies with decision timing: as previous studies^8,9^ indicate, stochastic sensory fluctuations, when conditioned on a choice, contribute to this bias. When diffusion governs WM dynamics, the distribution of memory states broadens over time; thus, more delayed decision timing leads to greater separation of memory states associated with different choices (colored dashed lines in **Figure 2A**), resulting in an increased decision-consistent bias (**Figure 2C**).

Next, suppose both diffusion and drift govern WM dynamics (**Figures 1B,2E**). Then, both biases vary with decision timing. A recent study on WM for color^26^ suggests that drift causes memory states to approach discrete attractors, leading to increased stimulus-specific bias over time, while many others^27–29,37–41^ attribute this bias to sensory encoding. Similar drift dynamics may also govern WM for orientation (gray arrows in **Figure 1B**), causing stimulus-specific bias to grow over time (gray vertical bars in **Figure 2F**). As previously noted, decision-consistent bias will also increase with decision timing due to WM diffusion (colored vertical bars in **Figure 2g**).

Furthermore, diffusion dynamics predict that the broadening of the estimate distribution in near-reference trials—indicating choice-induced bias—will be more pronounced in the late DM condition than in the early one (**Figures 2D,H**). This is because, despite equal choice-induced bias in both conditions (colored arrows in **Figures 2A,E**), its effect on distribution expansion intensifies with decision delay under diffusion dynamics (marginal distributions in **Figures 2A,E; Figure S1K**).

In summary, both diffusion-only and drift-diffusion models predict that, with decision delay, decision-consistent bias and estimation variability increase in near-reference trials (**Figures 2C,D,G,H**). However, they differ in stimulus-specific bias: it stays constant over time in the diffusion-only model but increases in the drift-diffusion model (**Figures 2B,F**).

### Growth and decision-timing dependency of behavioral biases

To determine whether stimulus-specific bias increases during the delay, we compared the bias magnitude at the early (4.5 s post-stimulus) and late (10.5 s post-stimulus) discrimination epochs. Since direct estimation errors were unavailable during discrimination, we inferred the bias from the psychometric curve for each stimulus orientation (**Figure 1F**), using the deviation of subjective equality from the actual orientation as a proxy. For this, we fitted the bias weight parameter, assuming the bias varies in magnitude but retains its shape (see **STAR Methods**). We found that the bias increased over time, with a significantly greater bias at the late compared to the early decision (**Figure 3A**).

**Figure 3.**
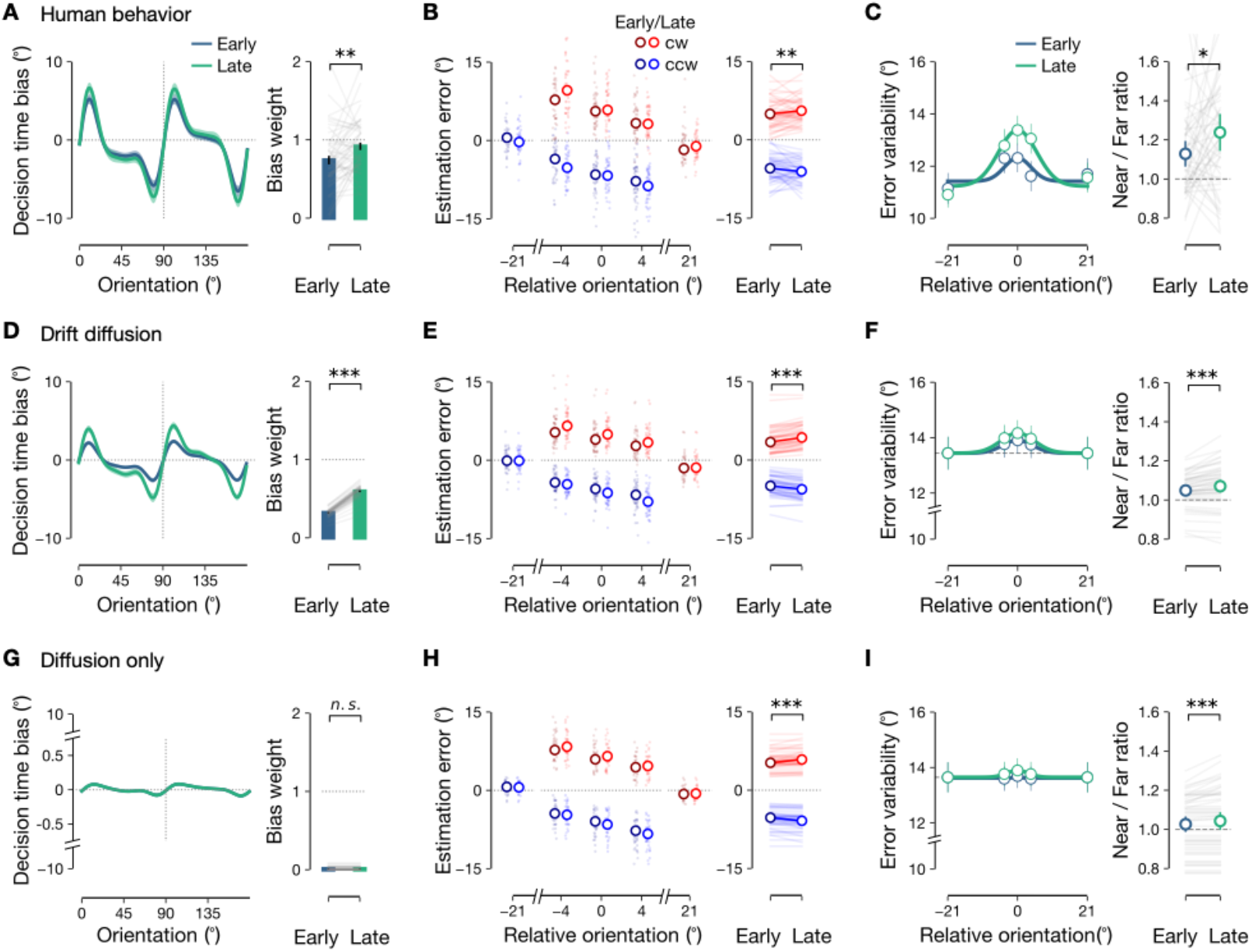
Biases in human behavior and models. **A-C**, Human data. **A**, Stimulus-specific bias at discrimination: Left, Bias estimates across orientations (lines with shades, means ± s.e.m.s across individuals); Right, Bias weights relative to estimation bias (bars, mean; lines, individuals; error bars, ± s.e.m; paired t-test, p = 0.0032). **B**, Decision-consistent bias in estimation (left) and their averages across near-reference trials (right): dots and thin lines, individuals; circles, medians ± s.e.m.; paired t-test, p = 0.0052. **C**, Error variability for early and late DM (left) and near-/far-reference variability ratio (right): circles, across-individual means; lines, individuals; Wilcoxon signed-rank test, p = 0.0450. **D-I**, Biases simulated by drift-diffusion (**D-F**) and diffusion-only (**G-I**) models. Format as in **A-C.** Paired t-test, p < 10^−10^ (**D**), p < 10^−10^ (**E**), p = 0.3614 (**G**), p = 0.0001 (**H**); Wilcoxon signed-rank test, p < 10^−5^ (**F**), p = 0.0007 (**I**). ***, p < 0.001, **, p < 0.01, *, p < 0.05, n.s., p > 0.05.

In contrast to their distinct predictions for stimulus-specific bias, both models predicted similar patterns of decision-consistent bias and near-reference variability (**Figures 2C,D,G,H**). Consistent with these, both measures were greater in the late DM condition (**Figures 3B,C**). To test the prediction regarding decision-consistent bias, we compared the reported orientation between CW-choice and CCW-choice trials for the near-reference condition; the difference was significantly larger in the late than the early DM condition (**Figure 3B**). For near-reference variability, we examined the difference in error variance between choice-unconditioned near-reference and far-reference trials; the difference was also greater in the late DM condition (**Figure 3C**).

To quantify drift dynamics, we fitted the models—with and without drift—to behavioral data (see **STAR Methods**). The model with drift outperformed the one without drift, according to the Bayesian Information Criterion (BIC) in conjunction with cross-validated log likelihoods (**Figure S2A**). Drift rate parameters indicate memory states drift modestly (less than 1°/s in median; *w*_*K*_ in **Figure S2C**). In *ex-post* simulations, the drift model accurately reproduced the observed growth and shape of stimulus-specific bias (**Figure 3D; Figures S3A,B**), unlike the non-drift model (**Figure 3G; Figure S3C**). Both models captured the observed decision-consistent bias patterns and variability increases in near-reference trials **(Figures 3E,F,H,I**), but a model solely based on efficient coding, without drift or diffusion dynamics, failed to do so (**Figures S3D-F**).

In summary, incorporating drift-and-diffusion dynamics with transient shifts in the chosen direction into WM effectively explains the growth of stimulus-specific and decision-consistent biases observed in our task.

### Influence of stimulus-specific drift on decision-consistent bias before and after DM

Our analyses indicate that memory states consistently drift toward attractors, with decision-consistent bias increasing with decision timing. This implies that decision-consistent bias follows distinct time courses depending on the stimulus’s position relative to these attractors in orientation space. We will first formalize this implication using our earlier drift-diffusion model, then test it against human data.

This implication involves dividing decision-consistent bias into its pre-decision and post-decision components. The pre-decision bias (*b*_*pre*_) refers to the difference in mean between the choice-conditioned distributions of memory states present at the onset of discrimination, whereas the post-decision bias (*b*_*post*_) develops after discrimination until estimation (see **STAR Methods** and **Method S1.1** for definitions).

Under diffusion-only dynamics, *b*_*pre*_ is expected to be greater in the late DM condition than the early DM condition, due to increased separation between choice-conditioned distributions (Δ*b*_*pre*_ > 0; dark gray patch in **Figure 4A**). Conversely, *b*_*post*_ should remain constant in both conditions if choice-induced bias remains unchanged in magnitude, as assumed (Δ*b*_*post*_ = 0; light gray patch in **Figure 4A**). These predictions were confirmed by the *b*_*pre*_ and *b*_*post*_ values derived from the *ex-post* simulation of the diffusion-only model (**Figure 4D**).

**Figure 4.**
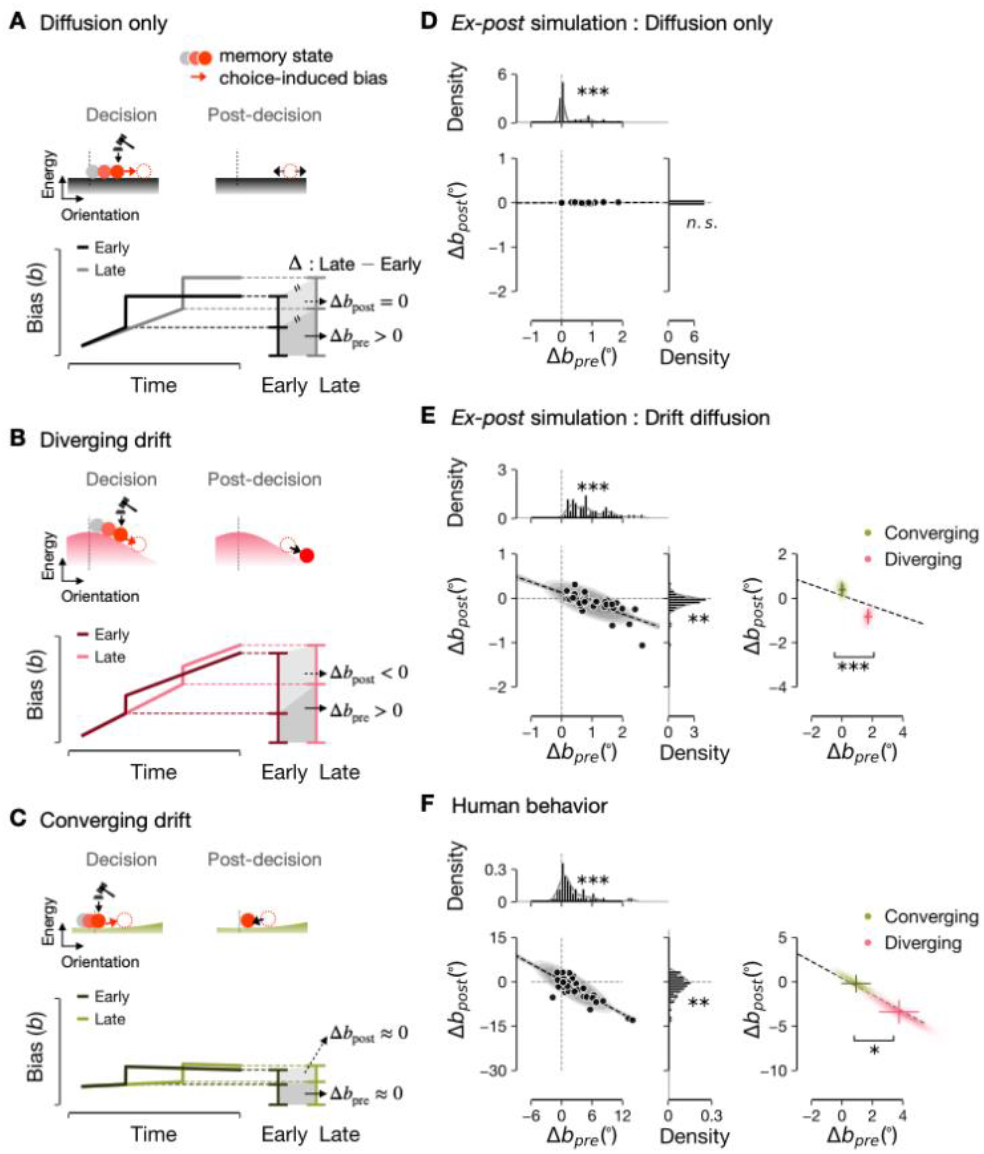
Decision-consistent biases before and after DM. **A-C**, Schematics of pre- and post-decision biases under diffusion-only (**A**), diverging-drift (**B**), and converging-drift (**C**) WM dynamics. Top, Memory dynamics on energy landscapes: gray to dark red circles, early to late states; dotted circles, states shifted by DM; red arrows, choice-induced biases; black arrows, post-decision biases. Bottom, Decision-consistent bias trajectories: dark/light lines, early/late-DM conditions. **D**, Across-individual Δb_pre_ and Δb_post_ values simulated by the diffusion-only model, shown as joint (left bottom) and marginal (top, right) distributions: dashed lines with shades, regression lines; dots, individuals. **E**, Simulations by the drift-diffusion model. Format as in **D**, except for the right panel, where across-individual averages of Δb_pre_ and Δb_post_ are shown separately for diverging and converging orientations. **F**, Human data, with format as in **E.** Signs of Δb_pre_ and Δb_post_ were tested with one-sample t-test (Δb_pre_, p < 10^−4^, Δb_post_, p = 0.5924 in **D**; Δb_pre_, p < 10^−10^, Δb_post_, p = 0.0024 in **E**; Δb_pre_, p < 10^−4^, Δb_post_, p = 0.0022 in **F**). Correlations were Pearson’s coefficients (r = 0.203, p = 0.1568 in **D**; r = −0.726, p < 10^−8^ in **E**; r = −0.829, p < 10^−10^ in **F**). Distances along regression lines were measured (1.941°, p < 10^−4^ in **E**; 4.215°, p = 0.0457 in **F**). *** p < 0.001, **, p < 0.01, *, p < 0.05, n.s., p > 0.05.

However, under drift-diffusion dynamics, the two components of decision-consistent bias display distinct, stimulus-dependent patterns. Consider the case where drift, diffusion, and choice-induced bias are moderate, so that memory states starting far from the attractors do not reach them during the delay, consistent with the best-fit parameters from the behavioral data (**Figure S2**). For orientations positioned between the attractors (e.g., cardinal orientations), memory states diverge from the stimulus (pink lines around 0° in **Figure 1B**), increasing the separation of choice-conditioned memory distributions over time, driven by the congruence between the diverging direction and the decision-consistent direction (**Figure 4B**). This leads to a greater *b*_*pre*_ in the late versus early DM condition (Δ*b*_*pre*_ > 0; dark gray patch in **Figure 4B**), while *b*_*post*_ should be smaller in the late DM condition (Δ*b*_*post*_ < 0; light gray patch in **Figure 4B**). Conversely, near the attractors (e.g., oblique orientations), converging drift decreases the separation of choice-conditioned memory distributions over time, counteracting the choice-conditioned separation (**Figure 4C**).

The drift-diffusion model also implies a covariation across individuals. While the best-fit drift rates are moderate (maximum *w*_*K*_ < 2.5°/s in **Figure S2C**), their differences across individuals predict systematic changes in decision-timing-dependent biases. Specifically, higher drift rates lead to more pronounced changes in both Δ*b*_*pre*_ and Δ*b*_*post*_. Thus, given their opposite signs and dependence on drift rate, a negative correlation between Δ*b*_*pre*_ and Δ*b*_*post*_ is predicted across individuals, driven by drift rate variability (**Figures S3G,H**).

These two implications were confirmed by the *ex-post* simulation data from the drift-diffusion model (**Figure 4E)**, along with human data analyses that did not rely on model estimates (**Figure 4F**). First, the Δ*b*_*pre*_ and Δ*b*_*post*_ were positive and negative, respectively, for diverging drift orientations (pink in right panels), but both were near zero for converging drift orientations (green in right panels). The joint distributions of Δ*b*_*pre*_ and Δ*b*_*post*_ were significantly separated. Here, considering the individual differences in stimulus-specific bias shape (**Figure 1G**), we determined whether a converging or diverging drift governs a given orientation based on participant-specific bias curves (see **STAR Methods**). Second, when the data were pooled across orientations (**Figures 4E,F**, left panels), the Δ*b*_*pre*_ and Δ*b*_*post*_ were still positive and negative, respectively. This was anticipated because the Δ*b*_*pre*_ and Δ*b*_*post*_ were large near diverging drift orientations. Across individuals, the Δ*b*_*pre*_ and Δ*b*_*post*_ were negatively correlated (**Figure 4F**).

In summary, phenomenological models reveal how the drift-diffusion dynamics of WM intricately shape decision-consistent bias before and after decisions in a stimulus-specific way, supported by human behavior.

### Drift-diffusion dynamics in cortical signals of orientation memory

The behavioral analysis examined the biases at snapshot moments of discrimination and estimation. To verify and expand on these findings beyond these moments, we decoded the WM signal of stimulus orientation from the blood-oxygenation-level-dependent (BOLD) measurements via inverted encoding analysis^42–44^, tracking the biases over time in that decoded signal. Focus was primarily on early visual areas, V1, V2, and V3, given their high-fidelity WM for orientation^43,45,46^, with parietal and frontal areas included for comparison (**Figures S4E-I**).

As implied by drift-diffusion dynamics, the cortical signal of orientation memory confirmed the growth of stimulus-specific bias over time. Stimulus orientation in WM was decodable from the early visual cortex with significant fidelity across all trial time points (**Figure S4D**), unattributable to eye movement confounds (**Figure S5**). Its trajectories, conditioned on stimulus orientations, drifted away from cardinal and toward oblique orientations (**Figure 5A**). To quantify these attractor dynamics, we tracked bias strength using linear regression weights that related each time point’s bias to each individual’s behavioral stimulus-specific bias (thin gray curves in **Figure 1G**). The bias weight was initially low, consistent with the efficient coding framework^28,29,37^ (**Figure S7**), and increased to match those observed in the behavioral errors (**Figure 5B**). Additionally, representational similarity analyses^47^ and simulated population responses with heterogeneous tuning curves (**Figure S6**; **Method S2.1**) confirmed the growth of stimulus-specific bias in memory representations.

**Figure 5.**
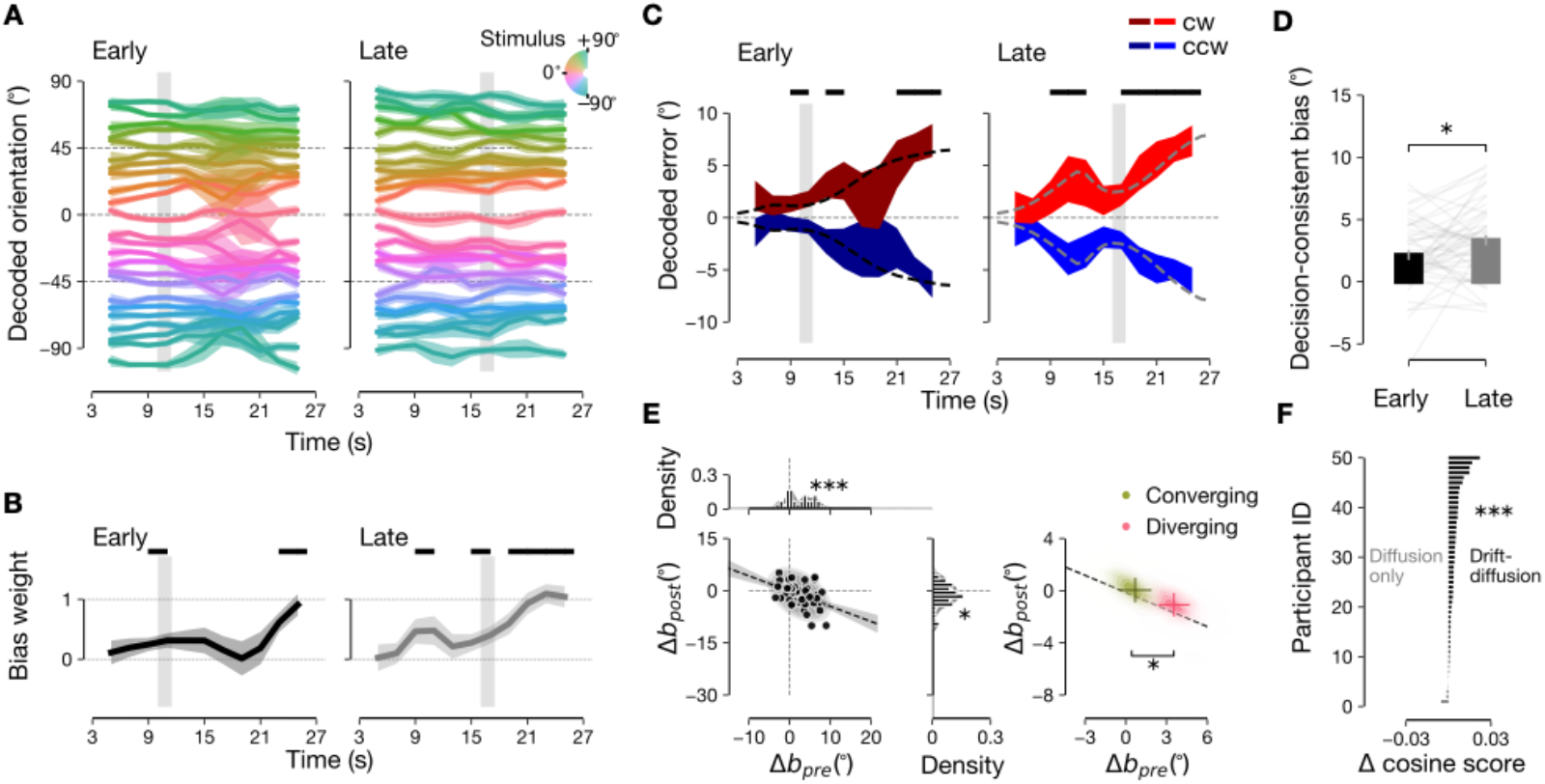
Cortical signals of stimulus-specific drift. **A,B** Evolution of stimulus-specific bias, depicted by decoded orientations (**A**) and bias weight relative to behavioral estimation bias (**B**). **C**, Evolution of decision-consistent bias, depicted by choice-conditioned decoded errors. Dashed lines mark the BOLD dynamics constructed from the piecewise linear fit (see **STAR Methods**). **A**-**C**, Gray bars indicate decision timing, with 4-s hemodynamic delay. Shades, ± s.e.m.s across trials. **B,C**, Black bars mark significant non-zero bias points (**B**) or between-condition differences (**C**): p < 0.05, permutation test, Bonferroni-corrected for time points. **D**, Decision-consistent biases at early and late DM, estimated from the model fit described in **C**: gray lines, individuals (p = 0.0176, paired t-test). **E**, Decision-timing-dependent changes in the pre-decision and post-decision biases, estimated from the model fit described in **C** (format as in **Figure 4F**; one-sample t-test, Δb_pre_, p < 10^−5^; Δb_post_, p = 0.0142; correlation between Δb_pre_ and Δb_post_, r = −0.431, p = 0.0018; distance along regression line, 3.069°, p = 0.0451). **F**, Model comparison in cosine scores of decoded cortical signals (paired t-test, p < 10^−6^). *** p < 0.001, * p < 0.05.

We note two caveats in inferring WM dynamics from BOLD signals. First, BOLD signals may appear to change more gradually than actual neural activity due to hemodynamic effects. Second, the limited temporal resolution of BOLD signals can cause interference between stimulus and reference orientations during the discrimination epoch. Indeed, the brain signals transiently shifted toward the near-reference orientation, especially in the late DM condition (**Figure 5A**, right), resulting in a transient drop in stimulus-specific bias (**Figure 5B**, right) around the discrimination epoch. To properly compare BOLD signals with model predictions, we addressed these issues through event-related anaylsis^48^: convolving the predicted trajectories of WM with the canonical hemodynamic response function (HRF), and incorporating memory attraction to reference orientation into the model prediction through regression weights (see **STAR Methods**). With these corrections, we examined whether cortical signals align with the implications of the drift-diffusion dynamics on decision-consistent bias.

The decision-consistent bias in BOLD signals was estimated as follows: (i) mean decoding errors conditioned on choice at each time point were calculated for near-reference trials (**Figure 5C**, colored shades); (ii) piecewise linear functions were fitted to these error trajectories (**Figure 5C**, dashed lines); and (iii) the bias was quantified by averaging the deviations of the fitted functions from zero at the estimation epoch. Consistent with diffusion-only and drift-diffusion dynamics, we found that the bias increased significantly with decision timing (**Figure 5D**). Importantly, the decision-timing-dependent measure of bias increase derived from the inferred trajectories (Δ*b*_*pre*_ and Δ*b*_*post*_) matched the predictions from the drift-diffusion model (**Figure 5E**), showing positive Δ*b*_*pre*_, negative Δ*b*_*post*_, negative correlation between Δ*b*_*pre*_ and Δ*b*_*post*_, and distribution separation between Δ*b*_*pre*_ and Δ*b*_*post*_ for the stimulus orientations with converging and diverging drifts.

Lastly, to assess how accurately cortical signals reflect behavioral biases, we convolved the predicted memory trajectories from the best-fit diffusion-only and drift-diffusion models with the canonical HRF (see **STAR Methods** and **Figures S4A,B**). The stimulus-specific trajectories predicted by the drift-diffusion model (**Figures S7A,B**) closely matched the BOLD signals of orientation memory (**Figures 5A,B**), unlike those from the diffusion-only model (**Figures S7C,D**). For most individuals, the cosine score indicated a better fit of the drift-diffusion model to the observed BOLD trajectories (**Figure 5F**).

In summary, the WM signals in the visual cortex corroborated the behavioral findings, confirming the implications of the drift-diffusion dynamics.

### RNNs with drift dynamics reproduce the human data

We demonstrated that both behavioral and cortical responses support the importance of drift dynamics in explaining the temporal evolution of biases, deriving their implications from phenomenological models. We utilized task-optimized RNNs^4,49,50^ to investigate whether simple network mechanisms could underlie WM and DM interaction beyond the phenomenological level.

We trained 50 independent RNNs using a task equivalent to that for humans and a joint loss function that penalizes both discrimination and estimation errors (**Figure 6A**). Given the spatial separation of the stimulus and reference, we fed these inputs to distinct RNN populations. The stimulus inputs were assumed to have greater variability near oblique compared to cardinal orientations, consistent with the efficient coding principle^27,36^. This heterogeneity in variability prompted the RNNs to drift toward oblique orientations, as this reduced the overall training loss.

**Figure 6.**
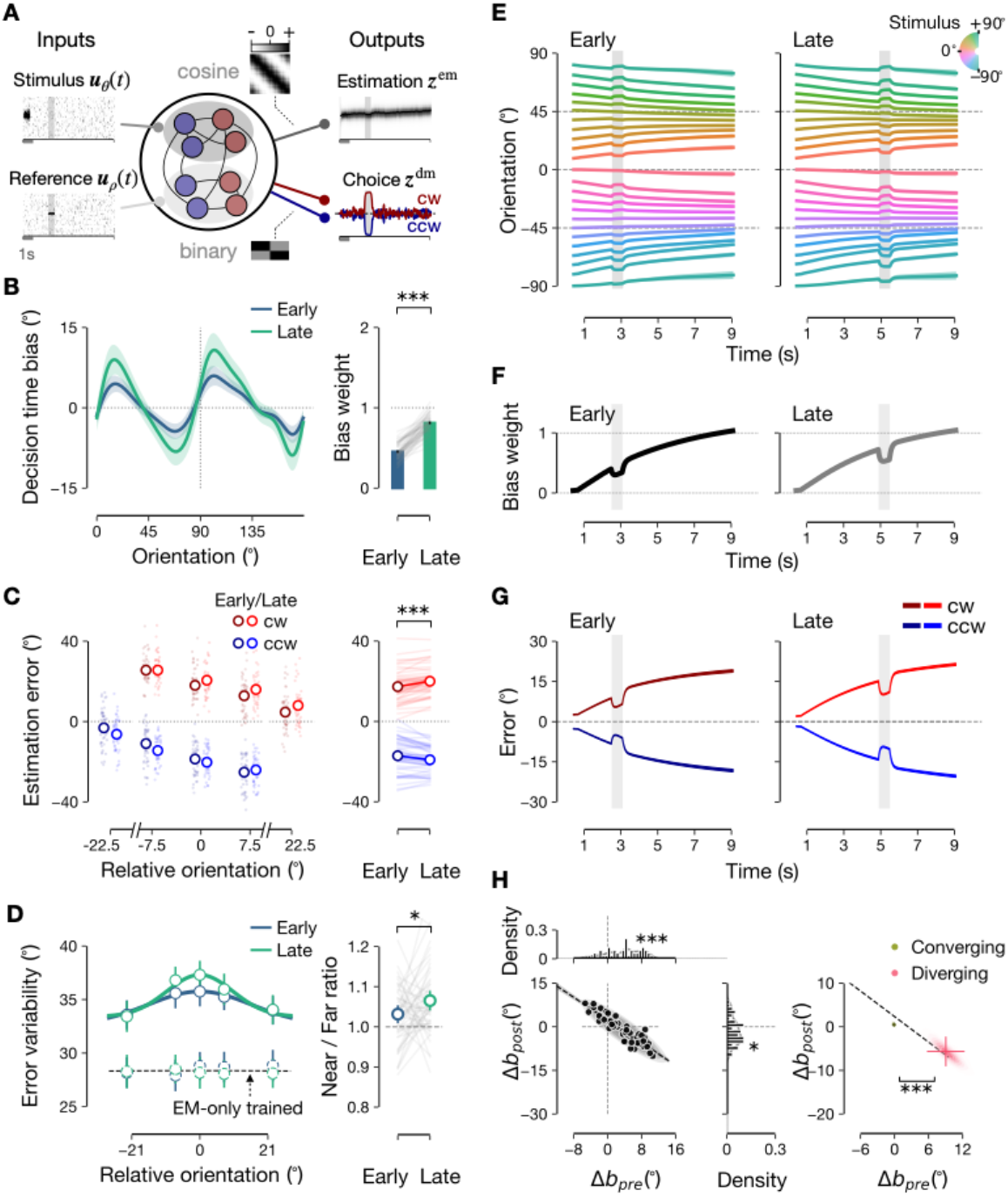
Biases in task-optimized RNNs. **A**, RNN architecture with time-varying inputs and outputs. **B-D**, Stimulus-specific bias growth (**B**), decision-timing-dependent decision-consistent biases (**c**), and near-reference variability (**D**). Format as in **Figures 3A-C.** In **D**, solid and dashed lines represent the original and estimation-loss-only RNNs. Paired t-test, p < 10^−10^ in **B**; p < 10^−10^ in **C**; Wilcoxon signed-rank test, p = 0.0390 in **D. E**-**G**, Evolution of stimulus-specific biases (**E,F**) and decision-consistent biases (**G**). Format as in **Figures 5A-C**, without hemodynamic convolution. **H**, Decision-timing-dependent changes in pre-decision and post-decision biases. Format as in **Figure 4F**. One-sample t-test, Δb_pre_, p < 10^−7^; Δb_post_, p = 0.0277; correlation between Δb_pre_ and Δb_post_, r = −0.897, p < 10^−10^; distances along regression line, 11.491°; p < 10^−4^. **, p < 0.01, ***, p < 0.001.

The trained RNNs exhibited all the features characteristic of human data, as implied by the drift-diffusion model. Stimulus-specific bias increased over time (**Figures 6B,E,F**), and decision-consistent bias grew (**Figures 6C,G**) with decision timing. A negative correlation was found between Δ*b*_*pos*t_ and Δ*b*_*pre*_ (**Figure 6H**, left), while Δ*b*_*pre*_ and Δ*b*_*pos*t_ were amplified in orientations with diverging drift (**Figure 6H**, right).

The RNNs displayed sensory drive effects linked to the reference presentation during the discrimination epoch (transient dips in **Figures 6E-G**), as assumed in our BOLD-response model. This is because all neurons in the RNNs, including those receiving the reference, contribute to the estimation (**Figure 6A**). The RNNs also exhibited increased variability in estimation errors in near-reference trials, indicating choice-induced bias, which increased with decision timing (**Figure 6D**, solid symbols). Importantly, these increases in estimation errors in near-reference trials vanished when RNNs were penalized only for estimation errors (**Figure 6D**, dashed symbols), highlighting DM’s critical role in generating choice-induced bias.

In summary, training RNNs to minimize both discrimination and estimation errors, while imposing drift dynamics, is sufficient for RNNs to display the main features of the estimation biases seen in human data.

### RNN mechanism for decision formation and choice-induced bias

The task-optimized RNNs exhibit a characteristic of choice-induced bias (**Figure 6D**), mirroring human behavior (**Figures 1I, 3C, S1A,B**). Investigating the formation of DM from WM and its impact on WM in RNNs may reveal the neural mechanisms underlying this bias, which remains elusive.

To avoid complications from drift dynamics, we studied the interplay between DM and WM in *homogeneous RNNs* trained on inputs devoid of orientation-specific variability, which therefore do not display drift. The average connectivity matrix showed a block-wise structure reflecting separate input and output pathways (**Figure 7A**), leading us to analyze interactions among three subpopulations: units receiving stimulus input (***r***_θ_), those receiving reference input and favoring the CW 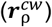 and CCW choices 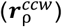 (**Figure 7B**).

**Figure 7.**
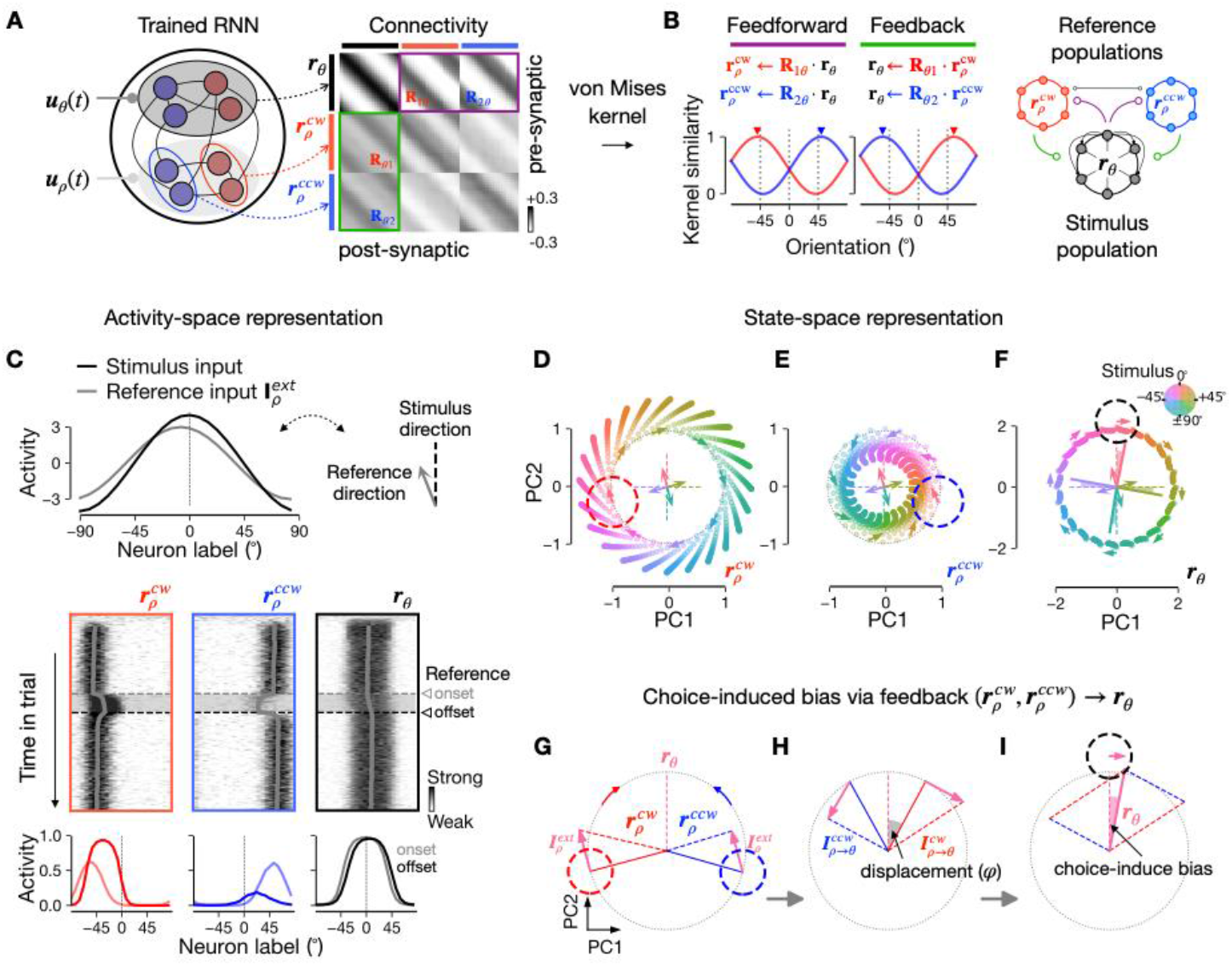
Mechanisms of decision-making and choice-induced bias in homogeneous RNNs. **A**, Subpopulations and average connectivities. **B**, Left, Scaled-rotation approximation of feedforward and feedback connections: **r**_θ_, stimulus-receiving units; 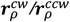, reference-receiving, CW/CCW-projecting units. Right, Three-ring system with rotation-based recurrent interactions. **C**, Activity changes during decision: Top, Input profiles over labeled neurons; Middle, Time courses of 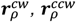, and **r**_θ_, with discrimination epoch marked with triangles; Bottom, activity snapshots at the onset (light) and offset (dark) of reference. **D-F**, Geometrical analysis of winning (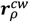, **D**) and losing (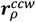, **E**) reference units and stimulus units (**r**_θ_, **F**) in 2D state space of **r** _θ_ during discrimination: isotropic rings, initial memory states; color saturation, time tracked for different stimulus orientations; short arrows, rotation directions; dashed lines and radial arrows, stimulus and reference input for four sample orientations. Dashed circles spotlight the rotation dynamics for 0° stimulus. **G-I**, Linear description of choice-induced bias using low-rank approximation of trained **J** (see **Method S4.1**). As 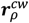 and 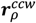 gravitate toward reference input 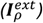, they shift outward and inward, respectively (**G**). Their feedback to **r**_θ_ rotates by a displacement φ (**H**), yielding summed inputs of 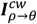 and 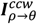 that bias activity in choice-consistent direction (black dashed circle in **I**).

All three populations exhibit bump-like activity through *feedforward* (***r***_θ_ to ***r***_ρ_) and *feedback* (***r***_ρ_ to ***r***_θ_) connections. The peaks of ***r***_θ_ encode and maintain stimulus orientations faithfully, while 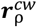 and 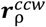 shift in *CCW* and *CW* directions, respectively (**Figure 7C**). These shifts reflect ***r***_θ_-to-***r***_ρ_ connectivity, well-approximated by scaled rotations in opposite directions (**Figure 7B**). Asymmetric connections and shifted memory representations resemble a head-direction system where interactions between opposing populations update direction in response to velocity signals^51^. Similarly, the input from ***r***_ρ_ updates ***r***_θ_ with a reference input (**Figure 7C**, shaded horizontal bars).

The feedforward dynamics from ***r***_θ_ to ***r***_ρ_ underlies DM. Suppose the stimulus is at 0° and reference is at −7.5°, placing the stimulus CW to the reference. Reference onset increases 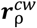, already rotated CCW, while 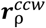 decreases (**Figure 7C**, bottom). The DM-mapping matrix reads this amplitude asymmetry into a choice that the stimulus is CW to the reference.

The DM mechanism can be analyzed geometrically through state space analysis (**Figures 7D-I**). In a 2D-PCA space defined by ***r***_θ_ (see **STAR Methods**), the memory manifolds of the three populations form a ring, initially with 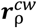 and 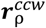 being rotated about 45° in the opposite directions from ***r***_θ_ (**Figures 7D-F**, dotted circles). During DM, reference input vectors (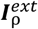, **Figure 7G**) are added to ***r***_ρ_, making 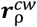 (winning population) expand and 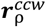 (losing population) contract (**Figures 7D,E**, arrows on circles). In geometrical terms, correct DM is achieved through a “rotation-addition” mechanism.

During DM, feedback dynamics implement choice-induced bias. Before DM, the feedback from 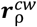 and 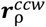 to ***r***_θ_ is balanced, keeping memory at the stimulus orientation. However, with 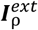 on, feedback from the winning population dominates, updating ***r***_θ_ in the choice-consistent direction, inducing bias (**Figure 7F**). After 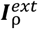 is off, ***r***_ρ_ quickly returns toward its pre-DM state (**Figures S8I,J**) and ceases reference-related influence. This supports our phenomenological model’s assumption (**Figures 1C,D; 2A,E**): choice induces an immediate, pulse-like update of memory states only during the discrimination epoch.

Further analysis revealed that choice-induced bias ultimately arises from a displacement in feedback dynamics. By linearizing the dynamics along the memory manifold, we found that the feedback rotation opposes the feedforward rotation but over-rotates with a displacement (denoted by *φ*; gray area in **Figure 7H** for over-rotation in 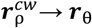). This creates an imbalance in feedback inputs 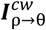 and 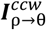 with 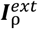 on, causing a choice-induced bias proportional to both 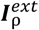 and *sin*(*φ*) (**Figures 7I, S8C-E**; **Method S4.2**). The essential role of the feedback connections is evident when they are ablated before training: discrimination remains intact, but the choice-induced bias does not emerge (**Figures S8K-M**). Furthermore, decision variable strength scales with choice-induced bias magnitude (**Figure S8F**), underscoring its functional significance in enabling robust DM under noise.

### RNN mechanism of the interaction between stimulus-specific and choice-induced biases

Building on our characterization of how the homogeneous RNNs instantiate choice-induced bias, we revisited the original heterogeneous RNNs—those exhibiting stimulus-specific bias—to identify the mechanism mediating the interaction between stimulus-specific and choice-induced biases. Despite differences in recurrent connectivity (**Figure S8B**), both RNNs represent stimulus orientations with ring manifolds in similar low-dimensional subspaces. This allows us to project the heterogeneous RNN responses onto the homogeneous RNN state space (**Figures 8A-C**).

**Figure 8.**
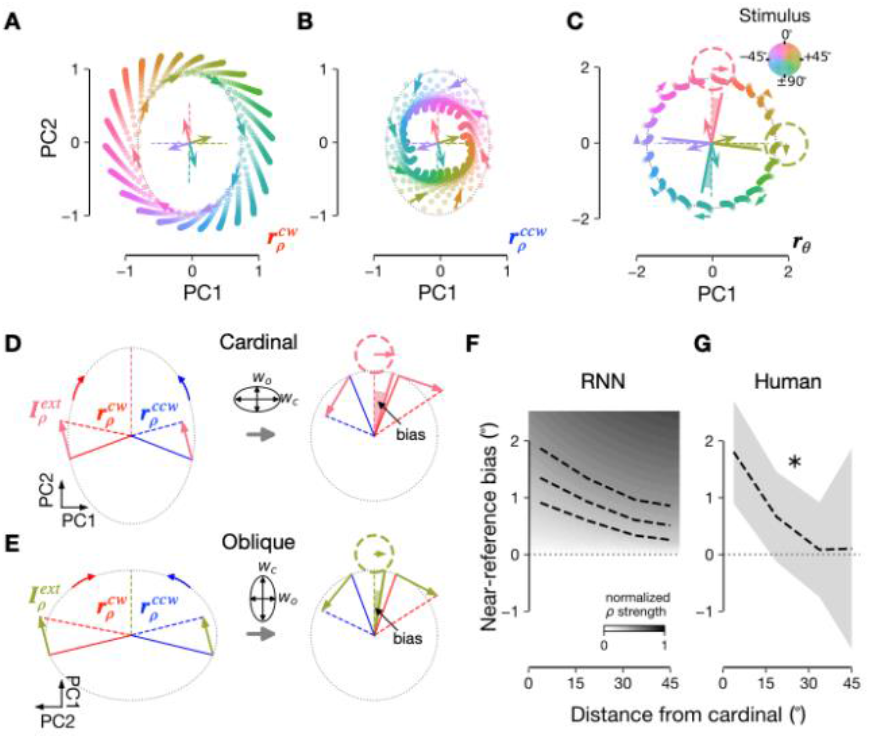
Orientation-dependent biases in heterogeneous RNNs. **A-C**, Drift-warped geometry of 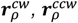, and **r**_θ_ during discrimination, shown in the same 2D state space and format as **Figures 7D-F. D,E**, Linear description of choice-induced bias for cardinal (**D**, 0°) and oblique (**E**, 45°) stimuli. The feedback from **r**_ρ_ to **r**_θ_ is anisotropic, producing greater elongation along cardinal (w_c_) than oblique (w_o_) orientations (ellipse above the gray arrows). With w_c_ > w_o_, choice-induced bias is more pronounced near cardinal orientations (right panels in **D,E**; see **Method S4.2**). **F**, Orientation dependency of near-reference bias. As reference signal increases in strength, near-reference bias increases while maintaining consistent orientation dependence (darker gray scales with three example bias patterns for different ρ values). **G**, Similar orientation-dependent bias observed in human behavior. One-sample t-test on individual slopes, p = 0.0304. *, p < 0.05.

In the heterogenous RNNs, ring manifolds in ***r***_*ρ*_ were warped into elliptic shapes before the reference input (**Figures 8A,B**), with cardinal orientations more sparsely represented than oblique ones, consistent with efficient coding theories for sensory network^32^. In drift dynamics, cardinal and oblique orientations correspond to diverging and converging stimuli, respectively. The warped geometry in ***r***_*ρ*_ effectively reverts to a circle shape in ***r***_θ_ by elongating along the minor axis more than the major axis (**Figure 8C**; **Method S4.1**). This anisotropic elongation causes 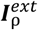 to have a stronger influence at diverging orientations (**Figure 8D**) than converging orientations (**Figure 8E**), resulting in greater choice-induced bias for diverging orientations (**Figures 8C,D**, dotted circles). Stimulus-specific drift further amplifies this effect: perturbations near diverging and converging stimuli are magnified and mitigated, respectively (**Figures 4B,C**), leading to larger biases for diverging orientations (**Figure 8F**). Human behavioral data validated this prediction: errors in near-reference trials were most biased around cardinal orientations and declined toward oblique orientations (**Figure 8G**).

In conclusion, the warping geometry of orientation representation and its anisotropic elongation are the key mechanisms mediating the intricate interplay of stimulus-specific, choice-induced, and decision-consistent biases in RNNs.

## Discussion

Two unique aspects of our paradigm enable us to identify decision-steered attractor dynamics as a source from which two crucial biases, stimulus-specific and decision-consistent biases, unfold interactively. First, the prolonged delay allows us to probe memory states at sufficiently distant moments through behavioral and neural measurements. Second, the mnemonic discrimination task eliminates sensory access to the target stimulus, and thus prevents decision processes from interfering with sensory encoding, unlike previous studies^2,8–10,12^. With this paradigm, we demonstrated that stimulus-specific bias intensifies over the delay while guiding decisions, and decision-consistent bias increases with decision timing in a stimulus-specific manner.

Analyzing the dynamics of task-optimized RNNs offered valuable insights into the WM-DM interactions. Simplification into three subpopulations, one for WM and two for DM, demonstrated how categorical decisions emerge from continuous orientation memory and how decisions immediately update memory, creating choice-induced bias. Central to both processes was asymmetric connectivity among these subpopulations, modeled as opposing rotation matrices, with their degree and scaling determining bias strength. This network property, within attractor dynamics, predicted stronger choice-induced bias for orientations with diverging drift, which was confirmed by behavioral data. Further, targeted ablation revealed that feedback from DM to EM populations, which causes the bias, can enhance decision robustness against noise, offering a rational basis for its presence.

Drift dynamics are not the sole source of stimulus-specific bias. Both our phenomenological and RNN models assume that sensory encoding variability—grounded in efficient encoding^27,28^—also contributes^36,37^. While downstream readout computations play a role as well^27–29,52^, the observed growth of stimulus-specific bias in behavioral, BOLD, and RNN data highlights ongoing WM updating via stimulus-specific drift during the delay. Notably, our use of “stimulus-specific drift” differs from “memory drift” in prior literature, which typically refers to random shifts in bump activity within a trial^14,53^. Many prior studies attribute such shifts to noise-driven diffusion^13,30^. In contrast, stimulus-specific drift refers to a systematic drift toward fixed attractors amid diffusion, evident when averaged across trials. For that matter, our BOLD-based analyses provide the first neural evidence of stimulus-specific bias growth through systematic drift over tens of seconds.

Stimulus-specific drift toward oblique orientations prompts questions about its mechanisms. Panichello et al.^26^ demonstrated that discrete attractor dynamics in delayed color estimation could reduce errors by biasing memory *toward* frequently encountered stimuli. However, as in our and prior studies^24,37^, orientation estimates are *repelled from* the frequent, cardinal orientations^27,28,36^ (similarly for location memory^33^). Thus, placing attractors around frequent stimuli does not apply to orientation. Instead, our phenomenological and RNN models propose that sensory input to the WM system varies according to the efficient encoding principle, explaining biases and variances inconsistent with traditional attractor models. Recent work^32^ indicates that orientation error evolution cannot be fully explained by single-module attractor models, emphasizing the roles of sensory and memory network interactions. Further work is needed to understand how these interactions influence error patterns across different features like color and orientation.

Our work clarifies source attributions for decision-consistent bias by differentiating drift toward attractors, stochastic noise^13,30^, and choice-induced bias^2,9^. Notably, stimulus-specific drift introduces a dynamic component: it initially biases memory, which biases choices, and then continues to bias memory in line with the biased choice. This has two key implications. First, models of DM and WM should account for how choices feed back into memory. Second, accurately explaining decision-consistent bias requires accounting for drift dynamics alongside stochastic noise, as neglecting this may misattribute post-decision drift to choice-induced bias. Additionally, it offers a perspective on confirmation bias^54–56^, suggesting it may arise from decision-consistent bias carried from pre-decision to post-decision phases, facilitated by stimulus-specific drift.

Beyond drift-diffusion dynamics, we provided a mechanistic account of choice-induced bias. Our phenomenological models assumed that decision formation transiently shifts memory states in the chosen direction during DM, supported by the RNNs. While this bias is attributed to the DM epoch, it may also originate earlier in encoding or later in decoding, with their contributions varying by task structure. Previous accounts^2,10^ suggested that choice-induced bias arises from a non-uniform weighting strategy optimized for DM and subsequently reused for estimation. This reliance on readout optimization makes it difficult to learn a stable decision boundary when reference inputs vary across trials. Consistent references may allow for late-stage readout strategies via selective information flow^57^ or memory recall^12^. In our paradigm, these optimization or selection strategies seem unlikely, as the reference varies each trial and is briefly available during DM. Further, unlike earlier proposals^9,12^, situating choice-induced bias during DM predicts specific neural trajectories, verifiable by examining the decision-related and memory-related neural responses with high temporal resolution, as shown by our RNNs.

Our work offers novel insights into how the brain processes task-relevant features before, during, and after categorical decisions, while optimizing performance in mnemonic discrimination and estimation. These insights warrant further validation and refinement. The mechanism by which our RNNs instantiate choice-induced bias—asymmetric connections and population dynamics—can be explored through synaptic connectivity and state-space dynamics^58^. Our integrated account of stimulus-specific and decision-consistent biases can also be extended to incorporate effects of memory load^24,59,60^ and serial dependence^61–63^, which may modulate WM dynamics, possibly through divisive normalization^37^ or short-term synaptic plasticity^64,65^. Overall, our work highlights the necessity of considering WM dynamics to fully understand perceptual biases with multiple origins, previously investigated in isolation.

## Acknowledgments

We thank Justin L. Gardner, Albert Compte, and Seth W. Egger for helpful comments. We thank Rosanne L. Rademaker, Matthias Fritsche, and Denis Schluppeck for making their data public. S. L. was supported by STI2030-Major Projects, No.2021ZD0203700/2021ZD0203705. S. L. acknowledges the support of the Shanghai Frontiers Science Center of Artificial Intelligence and Deep Learning and the NYU-ECNU Institute of Brain and Cognitive Science at NYU Shanghai. S.-H. L. was supported by the National Research Foundation of Korea (NRF) funded by the Ministry of Science and Information and Communications Technology (grant No. NRF-2021R1F1A1052020; NRF-2018R1A4A1025891; RS-2024-00349515), and by Korea Basic Science Institute (National research Facilities and Equipment Center) grant funded by the Ministry of Education (grant No. RS-2024-00435727).

## Author contributions

J.Lee., S.K., J.Lim, H.-J.L., M.J.C., D.-g.Y., and S.-H.L. designed research; H.G., S.L., and S.-H.L. performed research; H.G., H.-J.L., H.L., S.K., J.Lim and J.H.R. contributed unpublished reagents/analytic tools; H.G. wrote the first draft of the paper; H.G., S.L., and S.-H.L. edited and wrote the paper.

## STAR METHODS

## KEY RESOURCES TABLE

| REAGENT OR RESOURCE | SOURCE | IDENTIFIER |
| --- | --- | --- |
| **Deposited data** |  |  |
| Behavior data (preprocessed) | This paper | Open Science Framework:<br>https://osf.io/6q95m |
| fMRI data (preprocessed) | This paper | Open Science Framework:<br>https://osf.io/6q95m |
| Behavior and fMRI data (raw) | This paper | OpenNeuro:<br>https://openneuro.org/datasets/ds005381 |
| **Software and algorithms** |  |  |
| fMRIPrep 20.2.0 | Esteban et al. (2019)<sup>66</sup> | https://fmripiprep.org/en/20.2.0/index.html<br>RRID: SCR_016216 |
| MGL | Gardner et al. (2018)<sup>67</sup> | https://github.com/justingardner/mgl<br>doi: https://zenodo.org/records/1299497 |
| mrTools | Gardner et al. (2018)<sup>68</sup> | https://github.com/justingardner/mrTools<br>doi: https://zenodo.org/records/1299483 |
| Custom code | This paper | https://github.com/hyunwoogu/dynamic_bias |

## RESOURCE AVAILABILITY

## Lead contact

Further information and requests for resources and reagents should be directed to and will be fulfilled by the lead contact, Sang-Hun Lee at.

## Materials availability

This study did not produce new materials.

## Data and code availability

Raw behavior and fMRI data are publicly available at OpenNeuro: ds005381. Processed data are available at Open Science Framework: https://osf.io/6q95m. Original Python code for all analyses and figure generation is available at https://github.com/hyunwoogu/dynamic_bias.

## EXPERIMENTAL MODEL AND SUBJECT DETAILS

This study was conducted in accordance with the guidelines of and under the approval of the Institutional Review Board of Seoul National University. 50 healthy individuals (30 females, 19 − 32 years old; normal or corrected-to-normal vision) completed at least two or all of the three-day sessions of the main fMRI experiment across three days. Each participant provided written informed consent prior to the experiment and was naïve to the purpose of the study.

## METHODS DETAILS

### Experiments

#### Experiment stimuli and procedure

Stimuli were generated using MGL^67^ and presented by an LCD projector (60Hz). Participants viewed the stimuli at a visual angle of 22° (width) × 17°(height). The stimuli were displayed within a whole field (radius, 8.5°) gray circular Gaussian envelope aperture on a black background. A black fixation dot (0.07°) and a surrounding black fixation ring (0.83°to 0.9°) were constantly present at the center of the screen. There were 20 runs across the three scanning sessions, each run consisting of 12 trials. Some runs were excluded due to excessive motion (maximum motion across axes in rotation and translation surpassed T2^*^-weighted voxel size). Participants completed a one-hour practice session a few days before the main fMRI experiment.

Inside the scanner, participants maintained central fixation throughout each run and responded using a button box with linearly aligned keys labeled Key1 to Key4. Before each trial, the fixation dot dilated (0.14°) for 0.5s to cue the stimulus onset. Each trial started with a 1.5s presentation of an alternating (8/3 Hz) donut-shaped oriented grating (spatial frequency, 1 cycle/degree) spanning the peripheral visual field (aperture radii: inner, 2°; outer, 8.5°). The stimulus orientations ranged from 0° to 172.5° with a step size of 7.5°. The target stimulus presentation was followed by a first-epoch delay, a discrimination task, a second-epoch delay, and an estimation task. Two conditions were considered: early DM trials with 4.5s first-epoch and 10.5s second-epoch delays, and late DM trials with 10.5s first-epoch and 4.5s second-epoch delays.

In the discrimination task, participants viewed an oriented reference frame, a virtual line connecting the two yellow nonius dots (mark size, 0.1°) on the fixation ring. The fixation dot turned yellow and transiently dilated to 0.14° for 0.5s to cue the discrimination task onset. Participants indicated whether the target was tilted counter-clockwise or clockwise relative to the reference by pressing the Key2 (CCW) or Key3 (CW) with their left or right thumb, respectively. Relative orientations of the reference to the target stimulus were uniformly selected from [−21°, −4°, 0°, 4°, 21°], whose range approximately matches those of the previous studies^2,9^. Participants had 1.5s to respond, with their responses recorded without their knowledge, with a 0.5s buffer. If a response was made, the fixation dot dilated to 0.14° and turned blue for 0.75s; if no response was made within 1.5s, the fixation dot dilated to 0.14° and turned red for 0.75s. The reference frame disappeared upon button press.

In the estimation task, the participants reproduced the target stimulus from memory by rotating the two green nonius dots with Key2 (CCW) and Key3 (CW) within the 4.5s. The fixation dot turned green and transiently dilated to eccentricity 0.14° for 0.5s to cue the estimation task onset. Participants confirmed their adjustments by pressing Key1 using their thumbs. If a response was made, the fixation dot dilated to 0.14° and turned blue for 0.75s; if no response was made within 4.5s, the fixation dot dilated to 0.14° and turned red for 0.75s. The starting orientation of the estimation nonius dots was randomly chosen from 0° to 180°. The estimation task was followed by a 5.5s inter-trial interval (ITI). Each trial lasted 28s, with a total run time of 336s. After each run, participants received a summary of their performance.

#### MRI data acquisition and preprocessing

MR data were collected using a Siemens 3 Tesla Tim Trio with a 32-channel head matrix coil at the Seoul National University Brain Imaging Center. Participants underwent T1-weighted, high-resolution (0.8 × 0.8 × 0.8mm^3^) anatomical scans (repetition time (TR), 2.4*s*; inversion time (TI), 1s; time to echo (TE), 2.19ms; flip angle (FA), 8°). Over three separate days, they participated in the main T2*-weight fMRI scanning sessions: Day 1 with of retinotopy-mapping run (96s), hemodynamic impulse response function (HIRF) estimation run (96s), and 6 task runs (336s), Day 2 with 8 task runs, and Day 3 with 6 task runs. Scan parameters for the retinotopy-mapping, HIRF, and task runs were: voxel size, 2.3 × 2.3 × 2.3mm^3^; TR, 2.0s; TE, 30ms; FA, 77°. After the acquisition of fMRI scanner data, the initial preprocessing steps for the anatomical and functional images followed the fMRIprep workflow^66^ (version 20.2.0) with field map-free distortion correction option (–use-syn-sdc). For control analyses, we included the projections onto the FSL MNI space (MNI152NLIN6Asym).

#### ROI definition and voxel selection

The V1, V2, and V3 were defined using standard traveling wave methods^69^. Two 15°-wide wedge bowties on the vertical and horizontal meridians served as stimuli as in our previous study^70^. To measure the voxel-wise signal-to-noise ratio (SNR), we used a checkerboard whole-field impulse (radius, 8°) at a 1/24Hz frequency in the HIRF scan. Using the retinotopy scan, subjects’ V1, V2, and V3 regions across the left and right hemispheres and across dorsal and ventral areas were defined and combined for BOLD analysis. Voxel-wise SNR was calculated as the stimulus frequency (1/24Hz) amplitude in the HIRF scan divided by the average amplitude of frequencies above the third harmonics, discarding voxels with SNRs under two, following our previous research^71^. For control analyses, we used the publicly available visual atlas for IPS^72^ as well as the frontal regions^73^, IFC and DLPFC. Each voxel’s time series was converted into percent signal change by dividing by its average over the entire time series. To minimize the artifacts, fMRIprep-derived confounding variables were regressed out, consisting of white matter, CSF, and six additional three-dimensional motion regressors, along with the discrete cosine transform bases below 0.008Hz to reduce low-frequency components. No additional spatial smoothing was applied. The confounders were regressed out simultaneously to minimize the potential artifacts from the stepwise regression^74^. Resulting time series were z-scored voxel-by-voxel and run-by-run for further analyses.

### Analysis of data

#### Quantifying the stimulus-specific bias from behavior data

The stimulus-specific bias was quantified from the estimation data or the discrimination data. As for the estimation data, we computed the stimulus-conditioned means of estimation errors *ε*(*θ*), the difference between the estimation 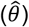 and the stimulus (*θ*), and fitted a smooth function *κ*(*θ*) to *ε*(*θ*). For each participant, the best-fit smooth function 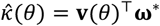 was found by finding ***ω*** that minimizes the sum of squared errors,

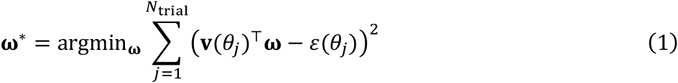

where 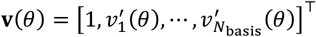 with 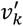 being the derivative of the von Mises density function with a center of 2*jπ*/*N*_basis_, a precision of *N*_basis_/2, and *N*_basis_ set to 12.

As for the discrimination data, we fitted psychometric functions *ψ* to the discrimination choices 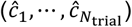 by maximizing the likelihood of choices ĉ ∈ {−1, +1}, corresponding to CCW (−1) and CW (+1), as follows:

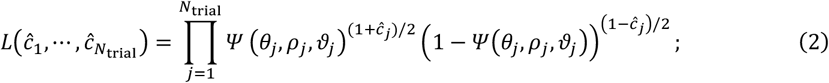

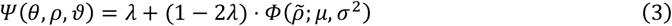

where *θ, ρ, ϑ* are stimulus orientation, reference orientation, and decision timing (*ϑ* ∈ {early,late}); 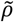 is the orientation of the reference relative to the stimulus 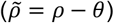; *Φ* is the cumulative Gaussian distribution function with a mean *μ* and a standard deviation *σ*; *λ* is the lapse rate. While fitting *ψ*, to parameterize the modulation of the stimulus-specific bias by decision timing (**Figure 3A**), we constrained the stimulus-specific mean of *Φ* with the best-fit smooth function 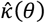, as follows: 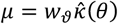, where, *w*_*ϑ*_ denotes the bias weight for the early (*w*_*early*_) or late (*w*_*late*_) DM conditions. Consequently, the maximum-likelihood fitting involved 5 free parameters in total: {*w*_*early*_, *w*_*late*_, *σ*_*early*_, *σ*_*late*_, *λ*}, where *σ*_*early*_ and *σ*_*late*_ are the standard deviations of *Φ* for early and late DM conditions, respectively.

To characterize the idiosyncratic patterns of stimulus-specific bias across participants, we defined the converging and diverging stimuli based on each individual’s 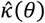. We first identified the zero-crossing of 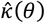 and estimated the local slopes. Then, for each participant, the nearby orientations (within ±8°) were labeled as diverging (if the slopes were positive) and converging (if the slopes were negative) stimuli.

#### Quantifying the near-reference variability from behavior data

As a signature of the choice-induced bias, the marginal distribution of estimation error spreads more in near-reference trials than in far-reference trials (**Figures S1D-K**). We referred to this signature as the *near-reference variability* and characterized it by comparing the variability of the reference-conditioned estimation error distributions across the reference conditions. As a robust measure of variability, we used the interquartile range (IQR), the difference between the first and third quartiles of the distribution. To capture the trend that the IQR increase as the reference nears the stimulus, we fitted a centered Gaussian density function, allowing the baseline, width, and amplitude parameters to vary.

To further validate our findings regarding the near-reference variability, we applied the same analysis procedure to publicly available datasets from previous studies (**Figures S1A-B**). In the work of Zamboni et al.^10^ and Fritsche & de Lange^8^, the reference was used as a decision boundary as in our study, allowing us to assess whether the near-reference variability is a generalizable signature of the choice-induced bias and is well captured by our procedure. Additionally, to further confirm that the near-reference variability does not occur when a decision-making is not imposed as a task demand, even in the presence of a reference-like stimulus, we also applied the same analysis to another publicly available dataset from Rademaker et al.^75^, where the intervening orientation stimulus acts only as a distractor (**Figure S1C**). For the Rademaker et al.’s dataset, we included a relative stimulus range of [−25°, 25°] in the analysis. Across all datasets, a reference was considered near if the relative stimulus orientation fell within [−8°, 8°].

#### Quantifying the decision-consistent bias from behavior data

To characterize the decision-consistent bias, we computed the conditional mean of estimation errors *ε* given choice *ĉ*, denoting it as *b* = (𝔼[*ε*|*ĉ* = *cw*] − 𝔼[*ε*|*ĉ* = *ccw*])/2. Previous studies^2,9^ showed that the decision-consistent bias is prominent only when the reference is near the stimulus orientation. Thus, we analyzed *b* only for the near-reference trials 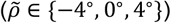 for further analyses.

#### Decomposition of the decision-consistent bias based on behavior data

We decomposed *b* into a component occurring before DM (pre-decision bias, *b*_pre_) and the one after DM (post-decision bias, *b*_post_):

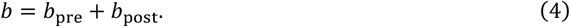

The decomposition was achieved in two steps, first quantifying the decision-consistent bias at the onset of DM from the discrimination data (*b*_pre_) and then quantifying the additional decision-consistent bias accumulated after DM up until the moment of estimation (*b*_post_). Conceptually, *b*_pre_ corresponds to the difference between the choice-conditioned means of memory states at the moment of discrimination time *t*_*dm*_,

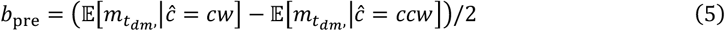

Here, the underlying distribution of memory states 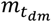, can be inferred from the parameters of the discrimination psychometric curve *ψ*(*θ, ρ, ϑ*) (defined in **Equation 3**), by applying the formalism offered by Signal Detection Theory^76^. Roughly put, this formalism relates the horizontal center and slope of *ψ* to the mean and dispersion of the inferred distribution of memory states 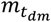. Then the choice-conditioned means of this distribution, 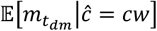 and 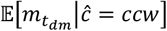, can readily be derived. We detailed this derivation in **Method S1.1**.

Next, having determined *b*_pre_, we quantified *b*_post_ from the distribution of estimation errors *ε*, as follows. First, to enable the single-trial estimation, we first sign-flipped the estimation errors according to the choice direction, aligning their signs with *b*_pre_, yielding sign-corrected errors *ε*^*^ = *ĉ* · *ε*. We then subtracted *b*_*pre*_ from *ε*^*^ for each trial to compute the residuals, *ε*^*^ − *b*_pre_. These residuals provide trial-to-trial estimates of how estimation errors are further deviated beyond *b*_*pre*_. To quantify the decision-timing dependent changes in *b*_post_, we performed a linear regression with condition indicators as regressors:

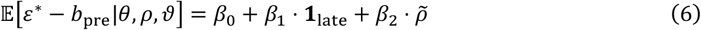

where *β*_0_ corresponds to the 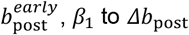 to *Δb*_post_, and *β*_2_ was introduced as a nuisance parameter to capture the previously known attraction towards the reference^75^.

For an additional comparison between the converging (conv) versus diverging (div) orientation conditions, we expanded **Equation 6** to incorporate the converging-vs-diverging orientation factor, as follows:

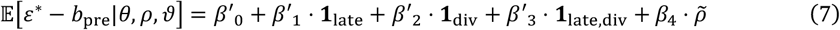

where *β*′_0_ corresponds to 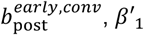 to 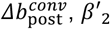 to 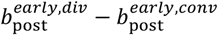 to 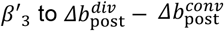, and *β*_4_ to the reference attraction.

#### BOLD decoding of orientation memory states based on inverted encoding analysis

We decoded stimulus orientation from the population BOLD responses in the early visual cortex (V1, V2, and V3) using inverted encoding analysis^42,43^. For each trial, the time courses of population BOLD responses **X** (*N*_voxels_ × *N*_trials_) were modeled as a linear combination of channel responses **Y (***N*_channels_ × *N*_trials_) with weights **W** (*N*_voxels_ × *N*_channels_), as follows:

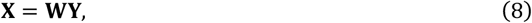

where each column of **Y** corresponds to the vector of channel responses to stimulus orientation *θ*_*j*_ in a given trial *j*: 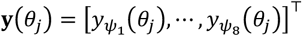, where *y_ψ_* (*θ*) = |cos(*θ* − *ψ*)|^8^ with channel centers *ψ*_1_, …, *ψ*_8_ tiling uniformly the orientation space [0, *π*].

In the following steps, we carried out the decoding analysis using a leave-one-run-out cross-validation procedure. First, we designated one run as a held-out *validation* run and the remaining runs as *train* runs. Second, we constructed a matrix of population BOLD responses **X**_T_ and a matrix of channel responses **Y**_T_ from the train runs, along with a matrix of population BOLD responses **X**_V_ from the held-out validation run. Third, given **X**_T_ and **Y**_T_, the weight matrix that yields the minimum squared errors **Ŵ** was determined as follows:

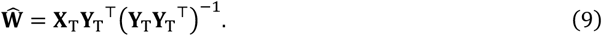

Fourth, we reconstructed the channel responses to the stimulus orientations presented in the validation run **Ŷ**_V_ based on **Ŵ** and the matrix of population BOLD responses in the validation run **X**_V_, as follows:

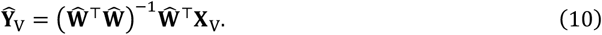

Fifth, to refine **Ŷ**_V_ at a fine scale, we repeated the third and fourth steps while repositioning the centers of the eight channels, thereby defining **Ŷ**_V_ for a total of 120 channel centers ***ψ*** = [0°, 1.5°, …, 178.5°]^⊤^. We repeated the whole steps for each run, decision timing, and time point, resulting in the reconstruction of channel response vectors **Ŷ**_V_(*t*), with the columns corresponding to all the trials for a given time point *t* of interest 3-14 TRs.

From these reconstructed channel responses **Ŷ**_V_(*t*), we decoded the single-trial stimulus orientation by mapping the reconstructed channel response in each trial *j* and time point *t* (**Ŷ**_*j*_(*t*)) to a point readout on the circular orientation space, as follows:

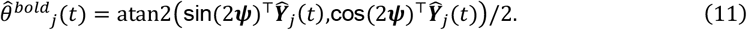

#### Estimating the time course of stimulus-specific bias in the memory states decoded from BOLD activity

The memory states decoded from BOLD activity exhibited the growth of the stimulus-specific bias over the delay. To track this growth, we estimated the amplitude of the bias at each time point based on the assumption that its across-stimulus profile is a scaled copy of the stimulus-specific bias estimated from the behavioral orientation estimates 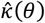, which was defined by the optimal parameters using **Equation 1** for each participant. Accordingly, for each participant and each time point of BOLD measurement, we fitted the multiplicative weight of 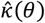 to the stimulus-specific errors of the memory states decoded from BOLD activity, 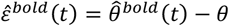. For further analyses, we also considered the sign-corrected decoding errors 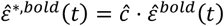, for which we flipped the signs of decoding errors 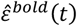 for each trial according to the choice direction.

#### Decomposition of the decision-consistent bias based on BOLD activity

For the BOLD activity, we estimated the decision-consistent bias (*b*^*bold*^) and its pre-decision 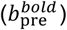 and post-decision 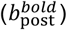 components in the following steps. First, we modelled the time course of the latent, decision-consistent bias in memory states with a piecewise linear function *g*(*t*) that incorporates the linear increase of the decision-consistent bias over time *t* along with the pulse-like shift due to the choice-induced bias during DM *t*_dm_, as we assumed in our phenomenological models:

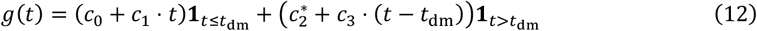

where the first term on the right-hand side of captures the initial bias by *c*_0_, the linear increase over time by *c*_1_ · *t* up to the time of DM, and the second term inherits the first term by including *c*_0_ and *c*_1_ · *t*_dm_ into 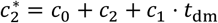 while capturing the pulse-like choice-induced bias by *c*_2_ and the linear increase over time by *c*_3_ · (*t* − *t*_dm_).

Second, we converted *g*(*t*) to 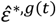 using a transfer function *v*, which convolves any given function with the canonical double-gamma hemodynamic response function^77^ *h*(*t*) while incorporating the input driven by the target stimulus *θ* and the reference stimulus *ρ*, which can be expressed as an argument of the complex number system as follows:

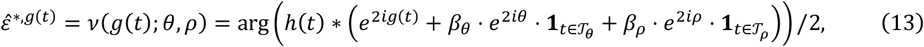

where *β*_*θ*_ and *β*_*ρ*_ denote the beta weights of visual events^48^ driven by the presentation of the stimulus *θ* and the reference *ρ*, while 𝒯_*θ*_ and 𝒯_*ρ*_ are the presentation time windows of *θ* and *ρ*.

Third, to estimate the influence of the stimulus and the reference (i.e., *β*_*θ*_ and *β*_*ρ*_), we assumed that the impact of DM is negligible in 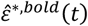 (as defined at the end of the previous section) in the far-reference trials and defined its model correspondence by plugging the zero bias in the transfer function defined in **Equation 13** instead of g(t): 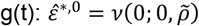 (see **Figure S4A**). Then, we found the values of *β*_*θ*_ and *β*_*ρ*_ that minimize the *L*_2_ difference between 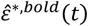 and 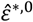.

Fourth, having estimated *β*_*θ*_ and *β*_*ρ*_, we then identified the parameters (*ĉ*_0_, *ĉ*_1_, *ĉ*_2_, *ĉ*_3_) of the time course of the latent, decision-consistent bias in memory states *g*(*t*) that minimize the L2 difference between 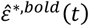 in the near-reference trials and 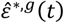.

Lastly, we obtained the estimates of the decision-consistent bias (*b*^*bold*^) and its pre-decision 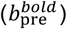 and post-decision 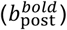 components, as follows:

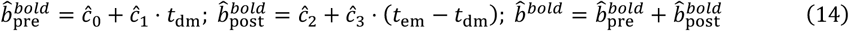

where *t*_em_ denotes the time of orientation estimation.

#### BOLD decoding of orientation memory states based on representational similarity analysis

As an alternative method for decoding orientation memory states from BOLD activity, we computed representational similarity matrices (RSMs) using the same population BOLD responses (**X)** used in the inverted encoding analysis. Following previous studies^78,79^, we averaged the across-voxel patterns of BOLD responses to each stimulus *θ* across trials (**x**_*θ*_) and then normalized **x**_*θ*_ by subtracting the across-stimuli mean 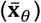 from it. We defined the correlation matrix **Ω** (*N*_stimulus_ × *N*_stimulus_) by computing the Pearson correlation of 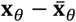 for each pair of orientation stimuli, while excluding identical cases that correspond to the diagonal elements. Such **Ω** was defined for each time point. We then estimated the orientation memory states 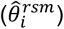 from **Ω** defined for each time point by taking the circular mean of each row, **Ω**_*i*_, corresponding to stimulus *θ*_*i*_:

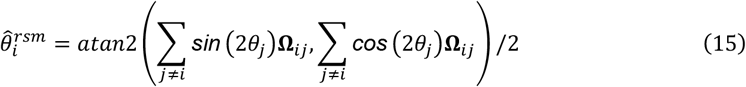

We quantified the “categorical” nature of orientation representation using a previously suggested category^40^: whether a stimulus is on the clockwise or counter-clockwise side with respect to the vertical orientation. To model this categorical pattern, we used a “block” matrix **M**_block_ (each row is +1 for within-category and −1 for across-category) as well as a “cosine” matrix **M**_cosine_ (each row is the cosine value of the corresponding stimuli). To quantify the relative contribution of each pattern, we used the weight *w*_cvx_ between zero and one, such that the convex combination **M**_cvx_ = *w*_cvx_**M**_block_ + (1 − *w*_cvx_)**M**_cosine_ approximates **Ω**, indicating a higher categorical representation by a higher value of *w*_cvx_. For comparison, we normalized **Ω** into the range [−1, +1] and used the least squares method to find *w*_cvx_ for each time point.

#### Eye-tracking

To ensure participants’ eyes remained fixed on the central fixation marker throughout the experiment, we monitored their eye positions using an MR-compatible video-based eye tracker (EyeLink-1000, SR Research). The eye tracker was set up at a sampling rate of 500 Hz. For each participant, we recalibrated the eye tracker before each session using the built-in five-point routine (HV5). Eye-tracking data were corrupted or not recorded for five participants due to technical issues. Data was further excluded from analysis for the scan runs where experimenters noted calibration issues, which were attributable to eye occlusion by the head coil, unreliable tracking due to reflective sources like MRI goggles, or excessive blinking patterns.

### Phenomenological models: diffusion-only and drift-diffusion models

#### Model description

We posited that the memory states in a single trial *m*(*t*) undergo the following dynamics within the orientation space spanning [0, *π*] with a periodic boundary:

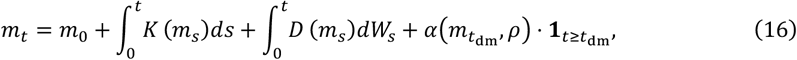

where *K*(*m*_*s*_) and *D*(*m*_*s*_) are the terms instantiating drift and diffusion dynamics, respectively, and *W*_*s*_ follows the Wiener process. We considered two classes of models, one with the diffusion term only (diffusion-only model) and the other with both drift and diffusion terms (diffusion-only model). The diffusion term *D*(*m*), which is shared by both models, was set to 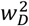. For the drift-diffusion model, the drift term *K*(*m*) was defined in a stimulus-specific manner to instantiate the stimulus-specific drift, as follows:

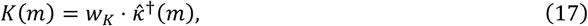

where *w*_*K*_ is the drift rate, and 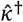 is the normalized stimulus-specific bias, 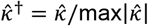, with 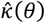 defined by the optimal parameters found using (**Equation 1**) for each individual.

The last term of the right side of **Equation 16** instantiates (i) the choice-induced bias by incurring an impulse-like shift in the memory trajectory in the choice-consistent direction and (ii) the reference-attraction bias at the moment of DM *t*_dm_:

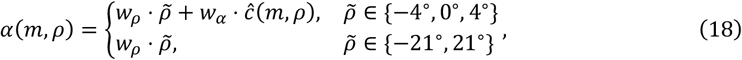

where 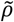 denotes the relative reference orientation; *w*_*ρ*_ is the strength of reference attraction, mimicking towards-distractor biases^75^; *w*_*α*_ is the strength of choice-induced bias only present in the near reference conditions 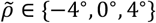, following the previous observations^2,9^. The choice term *ĉ* was determined by the relative difference between the reference orientation *ρ* to the memory state at the moment of DM: *ĉ* = *ĉ* (*m, ρ*) = sign(*m* − *ρ*).

To generate the discrimination and estimation reports, we used an instantaneous memory state at the corresponding moments of time: m(t) at *t*_dm_ = 6*s* and *t*_dm_ = 12*s* to determine *ĉ* in the early and late DM conditions, respectively; m(t) at *t*_em_ = 18*s* to determine an estimation report.

#### Constraining the initial memory states based on the principle of efficient sensory encoding

We constrained the stimulus-specific distributions of initial memory states *p*(*m*_0_|*θ*) based on the principle of efficient coding^27^. At the core of this principle is the encoding transformation function ℱ(*θ*), which captures how encoding resources are allocated. ℱ(*θ*) maps an orientation value *θ* in the stimulus space onto a measurement in the sensory space, in which the measurement is corrupted by the encoding noise. Therefore, using this framework, we first inferred ℱ(*θ*) from data and then used it to derive *p*(*m*_0_|*θ*).

We estimated the stimulus-to-sensory mapping ℱ based on a previously derived relationship between the stimulus-specific bias *κ*(*θ*) and the derivative of ℱ(*θ*)^28^:

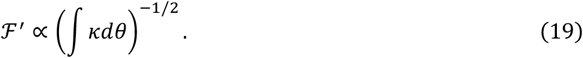

To compute the integral term in **Equation 19**, ∫ *κdθ*, we used previously estimated stimulus-specific bias 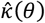 as an estimate for *κ*(*θ*). Given that 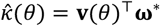, where 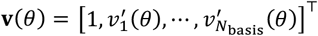 and ***ω***^*^ is obtained from **Equation 1**, the integral becomes **V**(*θ*)^⊤^***ω***^*^, where 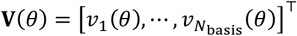 excluding the constant term. We then defined 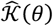, our estimate of ∫ *κdθ*, with an additional adjustment by a shape parameter *s* in [0,1], which controls the extent to which the stimulus-specific bias constrains ℱ^′^:

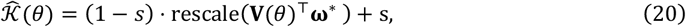

where rescale(·) denotes the min-max scaling between zero and one. As *s* approaches 0, the stimulus-to-sensory mapping becomes increasingly constrained by the integration of stimulus-specific bias function, and *s* = 1 corresponds to the case of uniform mapping. This estimator 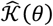 allows us to specify ℱ via **Equation 19**, by computing 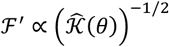 with the constraint ∫ ℱ^′^(*θ*^′^)*dθ*^′^ = *π* ^28^.

Having specified the encoding transformation ℱ(*θ*) and the sensory noise level *w*_*E*_, we can determine the initial distribution of memory states *p*(*m*_0_|*θ*) by modeling their initial states in the sensory space ℱ(*m*_0_) as a von Mises distribution centered around ℱ(*θ*) with dispersion proportional to *w*_*E*_. The density function is computable using the change of variables,

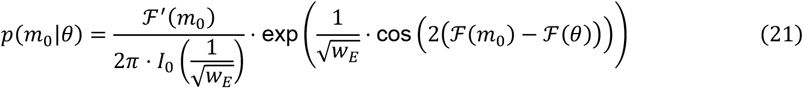

where ℱ^′^ is the derivative of ℱ, and *I*_0_ is the modified Bessel function of order 0. As such, we can fully constrain the stimulus-specific distribution of the initial memory states, *p*(*m*_0_|*θ*), with the previously estimated 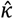 and additional two parameters, *s* and *w*_*E*_, which are fitted for both drift-diffusion and diffusion-only models (**Figure S2**).

#### Fitting the models to behavioral reports

To fit the models to the behavioral reports, we translated **Equation 16** into the corresponding Fokker-Planck equation:

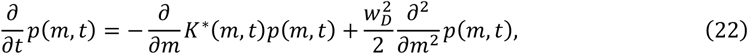

where *K*^*^(*m, t*) = *K*(*m*) + *α*(*m, ρ*)*δ*(*t* − *t*_dm_), where *δ*(·) is the Dirac delta function. We numerically solved the equation by discretizing *m* with a unit *Δm* = *π*/*N*_disc_, where *N*_disc_ = 96 (see **Method S3.1** for detailed numerical procedure for model fitting). For each participant, we fit the models to the discrimination choices *c*_*j*_ and the estimation reports 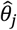 by finding the set of model parameters (8 parameters listed below) with the maximum joint likelihood *L*(parameters|data) given the experimental condition, which is specified by stimulus orientation *θ*_*j*_, reference orientation *ρ*_*j*_, and decision timing *ϑ*_*j*_:

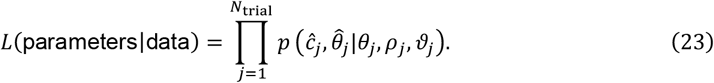

We used the optimization routines provided by SciPy, with 20 iterations, while randomizing initial parameters by drawing from the constrained ranges of the model parameters (see **Table S1**). The parameters set free to be fitted were *w*_*K*_ (drift rate) (set to 0 for the diffusion-only model), *w*_*D*_ (diffusion rate), *w*_*α*_ (choice-induced bias), *w*_*ρ*_ (reference bias weight), *w*_*E*_ (encoding variability), *w*_*P*_ (production variability), *w*_*λ*_ (decision-making lapse), and *s* (encoding function shape). To compute cross-validated log likelihoods, we ran 10 independent runs of 5-fold cross-validation of log likelihoods (each with 20 iterations) by separating the data used for fitting the models including the estimation of 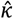.

#### Evaluating the correspondence between the drift-diffusion model and BOLD activity

We validated the drift-diffusion model’s prediction of the memory state dynamics by evaluating how closely it follows the trajectories of the memory states decoded from BOLD activity. For this evaluation, we translated the model prediction of memory states *m*_*t*_, as defined in **Equation 16**, into its equivalent in BOLD signal using the transfer function *ν*(*m*_*t*_; *θ, ρ*), as defined in **Equation 13**. Then, for each trial *j* in the near-reference conditions, we evaluated the correspondence between the model prediction *ν*(*m*_*j*_; *θ*_*j*_, *ρ*_*j*_) and the memory states decoded from BOLD activity 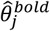 by quantifying the average cosine distance between them with a correspondence score 𝒮_*j*_, as follows:

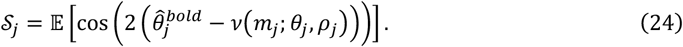

## Recurrent neural network model

### RNN dynamics

The following equations describe the dynamics of RNN:

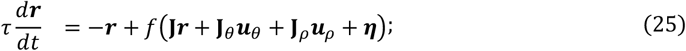

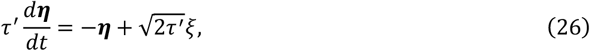

where ***r*** is the *N*_rec_-dimensional (*N*_rec_ = 96) unit activity with a time constant *τ* = 100 *ms*, and ***η*** is the stochastic noise with a time constant *τ*^′^ = 200 *ms*, modeled as the Ornstein–Uhlenbeck process^80^; ***u***_*θ*_ and ***u***_*ρ*_ are the *N*_in_-dimensional (*N*_in_ = 24) stimulus and reference inputs, respectively, whose units are orientation tuned, modeled as a von Mises distribution; **J, J**_*θ*_ and **J**_*ρ*_ are the weights of the recurrent, stimulus, and reference inputs, respectively; *f* = 1/(1 + exp(−*x*)) is the sigmoid activation function; *ξ* is the independent Gaussian noise with a standard deviation of 0.05. The values for ***r*** and ***η*** were initialized at 0s. We approximated the equations above using the forward Euler approximation with a discretization time step *Δt* = 20 ms.

Considering that the stimulus and reference inputs occupied different parts of the visual field in the task paradigm, ***u***_*θ*_ and ***u***_*ρ*_ were projected onto two separate 48-dimensional populations of the recurrent units ***r***, namely ***r***_*θ*_ and ***r***_*ρ*_. ***u***_*θ*_ is centered at veridical orientation *θ*, given as

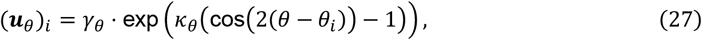

where *θ*_*i*_ is the preferred orientation of the unit *i*; *γ*_*θ*_ is the strength of the stimulus input, fixed at 1; *κ*_*θ*_ is the concentration parameter, fixed at 5. ***u***_*ρ*_ during the discrimination epoch was determined by a one-hot vector:

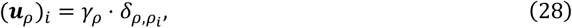

where *ρ* is the reference orientation, and *ρ*_*i*_ is the preferred orientation of the unit *i*; *γ*_*ρ*_ is the strength of the reference input, fixed at 2, which is higher than the one for ***u***_*θ*_ considering the higher level of certainty. We used *N*_*θ*_ = 24 ranging from 0° to 172.5° with 7.5° increments. The reference input was constrained to |*ρ* − *θ*| ≤ 30° with 7.5° steps, resulting in 9 possible relative references. During the “train episode,” we excluded *ρ* = *θ* to facilitate training but included it during the “generalization episode” (see the next section for the definition of the “train episode” and “generalization episode”).

Discrimination and estimation outputs, ***x***^*dm*^ and ***x***^*em*^, were

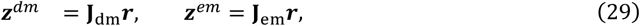

where 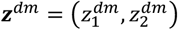 for CW and CCW choices, respectively, and ***x***^*em*^, a 24-dimensional ‘labeled line’ response vector, consists of equally discretized points within [0, *π*]. Input and output weights, **J**_*θ*_, **J**_*ρ*_, **J**_em_, and **J**_dm_, were fixed, defined as

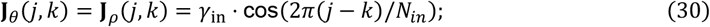

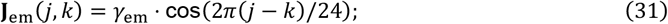

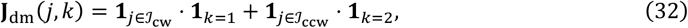

where *γ*_in_ = 1; *γ*_em_ = 0.4; a balanced voting for the two choices, with *𝒥*_cw_ = {*j* + 1: (*j* mod 48) 24} and *𝒥*_ccw_ = {*j* + 1: (*j* mod 48) ≥ 24}.

### RNN training

We trained RNNs using a paradigm equivalent in structure to the one used for human participants. RNNs were first trained on a short task timescale (“train episode”) and then generalized to an extended timescale (“generalization episode”). As we confine RNNs as a proof of principle, the time scales of dynamics were not directly aligned with the human experiment. Each trial in the train episode had the following structure: an initial fixation epoch with no inputs (0.1s) was followed by stimulus presentation (0.6s), first delay (0.3s), DM (0.6s), second delay (0.3s), and estimation report (0.1s) epochs. In the generalization episode, respecting the human task structure, the first and second delays were extended to 1.8 s and 4.2 s for the early DM condition, and 4.2 s and 1.8 s for the late DM condition, while the lengths of the other epochs remained the same.

For the supervised learning, we defined desired outputs ***q***^*dm*^ and ***q***^*em*^ as

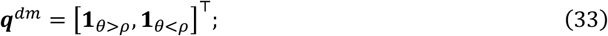

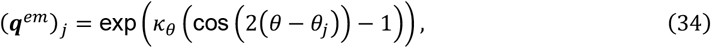

where ***q***^*dm*^ and ***q***^*em*^ were 2-dimensional and 24-dimensional vectors, respectively. We trained the recurrent weight **J** while maintaining other weights fixed. Before training, **J** was initialized as zero. The joint loss ℒ was ℒ_dm_ + ℒ_em_, where both ℒ_dm_ and ℒ_em_ are the time-averaging cross entropies between the network output and the desired output, given as:

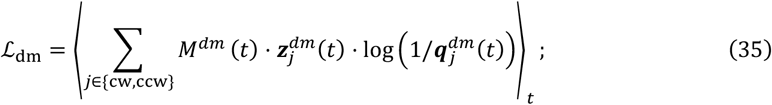

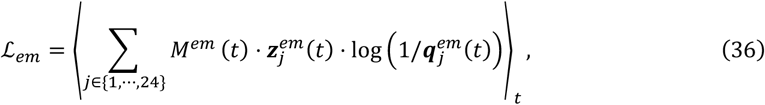

where *M*^*dm*^(*t*) is a binary mask, non-zero only during the discrimination epoch, while *M*^*em*^(*t*) is non-zero except for the initial fixation epoch. The loss was minimized using backpropagation in PyTorch with the Adam optimizer (with a learning rate of 0.02). We undertook 300 iterations per network training, generating 128 trials per iteration. In each of those trials, the stimulus and relative reference orientations were determined randomly.

To dissociate the effects of drift dynamics and the choice-induced bias, we independently trained 50 “homogeneous” RNNs, along with the original “heterogeneous” RNNs. For the heterogeneous RNNs (**Figures 6,8**), to approximate the orientation-specific variability that reflects the efficient sensory encoding principle, we added Gaussian noise to the orientation input *θ*, allowing the centers of ***u***_*θ*_ for a given stimulus orientation *θ*_0_ to vary as follows:

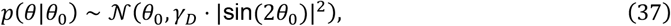

where the spread term *γ*_*D*_ was set at 10^2^. In contrast, no such noises were added for the homogeneous RNNs (*θ*|*θ*_0_ = *θ*_0_; **Figure 7**). All training details, except for stimulus input-level encoding variability, were identical for the homogeneous and heterogeneous RNNs.

To inspect the effect of feedback connections on the choice-induced bias, we independently trained 50 feedback-connection-ablated RNNs by zeroing the connectivity from the units receiving ***u***_*ρ*_ to those receiving ***r***_*θ*_. To examine the effect of fine-tuning the readout connection after training the feedback-ablated RNNs, we further trained the readout connection independently (mapping from recurrent activities ***r*** to both discrimination and estimation outputs ***z***^*dm*^ and ***z***^*em*^).

### RNN analysis

From the output vectors ***z***^*dm*^ and ***z***^*em*^ of the 50 independently trained RNNs, we determined their discrimination choice *ĉ*^*rnn*^ and estimation report 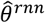, as follows:

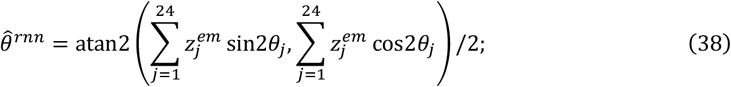

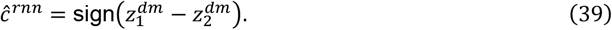

We then conducted the same analyses on *ĉ*^*rnn*^ and 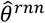, as we did on human discrimination choices *ĉ* and estimation reports 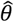, to assess the stimulus-specific and decision-consistent biases.

To depict RNNs’ population dynamics during the discrimination epoch in a low-dimensional state space, we applied PCA on average RNNs, taking the mean of individually trained ***J*** for the homogeneous and heterogeneous RNNs separately. We generated trials from both types of RNNs without network noise (*i*.*e*., *ξ* = 0 in **Equation 26**) for this state-space analysis. We analyzed the dynamics and connectivity patterns by further separating both ***r***_*θ*_ and ***r***_*ρ*_ into the CW- and CCW-projecting populations based on ***J***_dm_ (e.g., 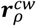 denotes the reference-receiving and CW-projecting population). For illustration, we used a near reference, *θ* − *ρ* = 7.5°, with winning/losing populations as ***r***^*cw*^/***r***^*ccw*^.

To inspect the dynamics of the homogeneous and heterogeneous RNNs, we projected the activities of mean homogeneous and heterogeneous RNNs onto the principal components from mean homogeneous RNNs, assuming slow drift dynamics in heterogeneous RNNs. We first stacked the activities of 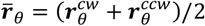 of the mean homogeneous RNNs for each condition (*N*_*θ*_ = 24 stimulus orientations and *N*_*ρ*_ = 9 relative reference orientations) to form a column-mean-centered data matrix 𝒟, that is, (*N*_*θ*_ · *N*_*ρ*_ · *T*) × 24, where *T* is the total time steps. In the text, ***r***_*θ*_ is used in the place of 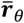 for brevity. The PC projection matrix **V**_𝒟_ was computed from the singular value decomposition of 𝒟 as 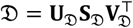. We used the first two PCs to project population activities of 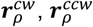, and 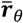. The first two PCs explained more than 92% of the total variance. For intuitive presentation, we sign-flipped and rotated the projection axes to align different stimulus conditions clockwise, with 0° stimulus points upright in ***r***_*θ*_ space, resulting in oblique stimuli (±45°) along the *x*-axis and cardinal stimuli (0°, 90°) along the *y*-axis.

## QUANTIFICATION AND STATISTICAL PROCEDURES

For the quantitative evaluations of phenomenological models (**Figures 4D,E, 5F**), we simulated the trajectories of memory states based on 10,000 Monte-Carlo iterations. For the statistical analyses of population-level differences in the decision-consistent biases around converging and diverging fixed points in the human behavior and drift-diffusion model (**Figures 4E,F**), the BOLD signals (**Figure 5E**), and RNN models (**Figure 6H**), we ran bootstrap-based permutation test using 10,000 random iterations.

## Supplemental Information

Document S1. Figures S1-S8, Methods S1-S4, Table S1, and supplemental references.

---

**Figure S1.**
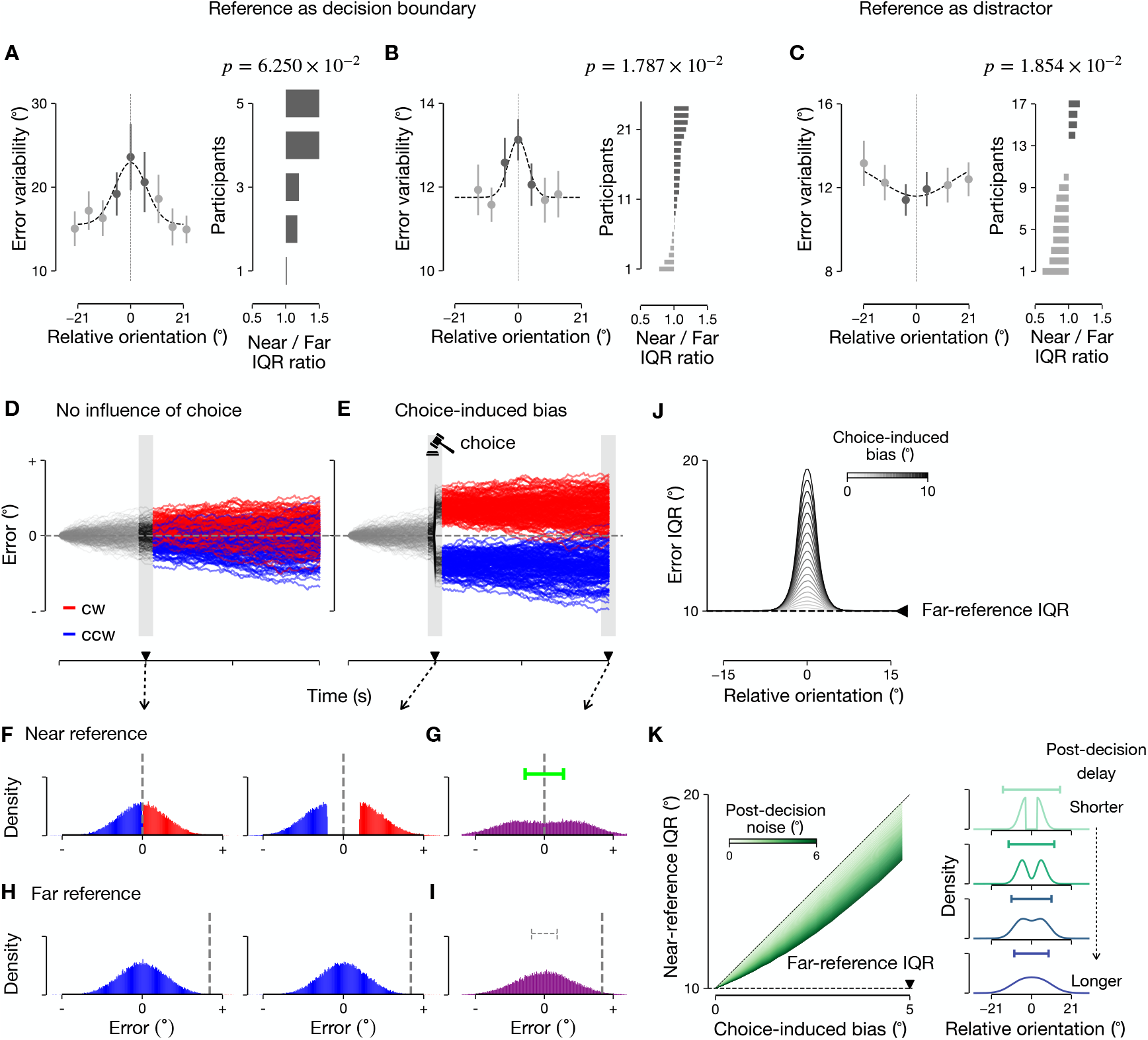
Behavioral signatures of choice-induced bias in marginal distribution of estimation errors, Related to Figures 1 and 2. **A-C**, Near-reference variability in datasets. **A,B**, Increased near-reference variability when the intervening reference serves as a boundary for decision, across the datasets of Zamboni *et al*. [S1] (**A**) and Fritsche & de Lange [S2] (**B**). In left panels, error variability, measured by interquartile range (IQR), is plotted as a function of stimulus orientation, relative to the reference orientation. Error bars denote ± s.e.m.s across participants. In right panels, the ratio of IQR in the near-reference condition compared to that in far-reference condition is plotted for each participant (sorted by their ratios). An IQR ratio larger than 1 indicates a wider error distribution in the near-reference condition. **C**, Decreased near-reference variability when the intervening reference serves as a distractor in the dataset of Rademaker *et al*. [S3] One-sample *t*-tests against 1, *t* (49) = 5.433 (*a*), *t* (4) = 2.701 (**A**), *t* (23) = 2.381 (**B**), *t* (16) = − 2.427 (**C**). Relative reference was considered “near” when the relative stimulus (*i*.*e*., stimulus minus reference) is within [−8°, −8°] across all the data sets (see **STAR Methods**). **D-E**, Effect of choice-induced bias in phenomenological models. Trajectories of choice-conditioned errors (red, CW choice; blue, CCW choice) under no influence of choice (**D**) and under choice-induced bias (**E**). Choice-induced bias is posited to be a transient jump that occurs at the moment of decision-making. **F**-**I**, Across-trial distribution of memory states, conditioned on choice at the time of discrimination (**F,H**) and unconditioned at the time of estimation (**G,I**). **J**, In simulation, the magnitude of near-reference variability increases in proportion to the size of choice-induced bias. **K**, Left, diffusive noise following the choice (greener lines with higher noise) dampens the magnitude of IQRs. When post-decision noise is absent (dashed diagonal line), the IQR difference from the far-to near-reference condition approaches twice the choice-induced bias. Right, Schematic depiction of the attenuation of near-reference variability in the marginal distribution as a function of post-decision delay.

**Figure S2.**
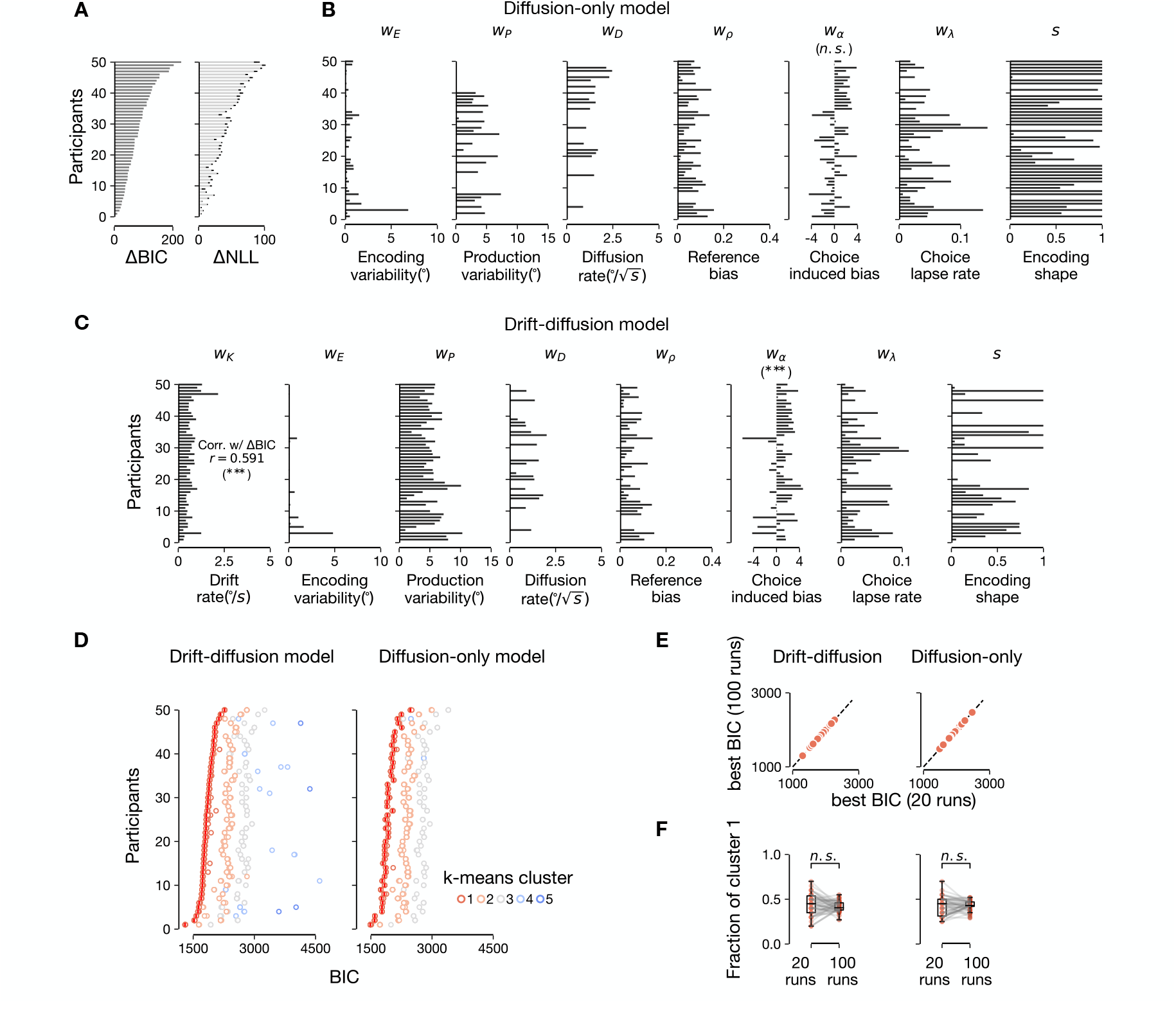
Goodness of fit of phenomenological models and parameter estimates, Related to Figure 3. **A**, Model comparison based on Bayesian information criterion (BIC, left) and cross-validated negative log likelihood (NLL, right, sorted by the values of ΔBIC) across participants. Positive values in ΔBIC and ΔNLL indicate the evidence in favor of the drift-diffusion model. Across all the participants and metrics, data suggested a superior fit for the drift-diffusion model. Lines denote ± s.e.m. **B,C**, Best-fit parameters of the diffusion-only model (**B**) and the drift-diffusion model (**C**). Participants were sorted according to their values of !BIC. Fitted choice-induced bias, *w*_*ω*_ in the diffusion-only model was not significantly different from 0 (Paired samples *t*-test, *t* (49) = 0.852, *p* = 0.398, 95% CI=[−0.359°, 0.888°]), while *w*_*ω*_ in the drift-diffusion model was significantly larger than 0 (Paired samples *t*-test, *t* (49) = 3.715, *p* = 5.218×10^−4^, 95% CI=[0.524°, 1.758°]). Fitted memory drift rate in the drift-diffusion model, *w*_*K*_, was correlated with the gain in BIC (ΔBIC) (Pearson correlation, *p* = 6.287×10^−6^, 95% CI=[0.374, 0.746]). **D**, BIC values from 20 independent model-fitting iterations (circles) for each participant and model, along with the best BIC values from additional 100 iterations (lines). Colors indicate score clusters identified using *k*-means clustering, where the number of clusters was determined based on silhouette scores. Participants were sorted according to their best BIC values for the drift-diffusion model. **E**, Comparison of the best BIC values obtained from the initial 20 iterations versus the additional 100 iterations. **F**, Proportion of BIC scores assigned to the *k*-means cluster (which includes the best BIC values) obtained from the initial 20 iterations versus those from the additional 100 iterations, based on the same clustering parameters. ^***^ *p <* 0.001, ns *p* > 0.05.

**Figure S3.**
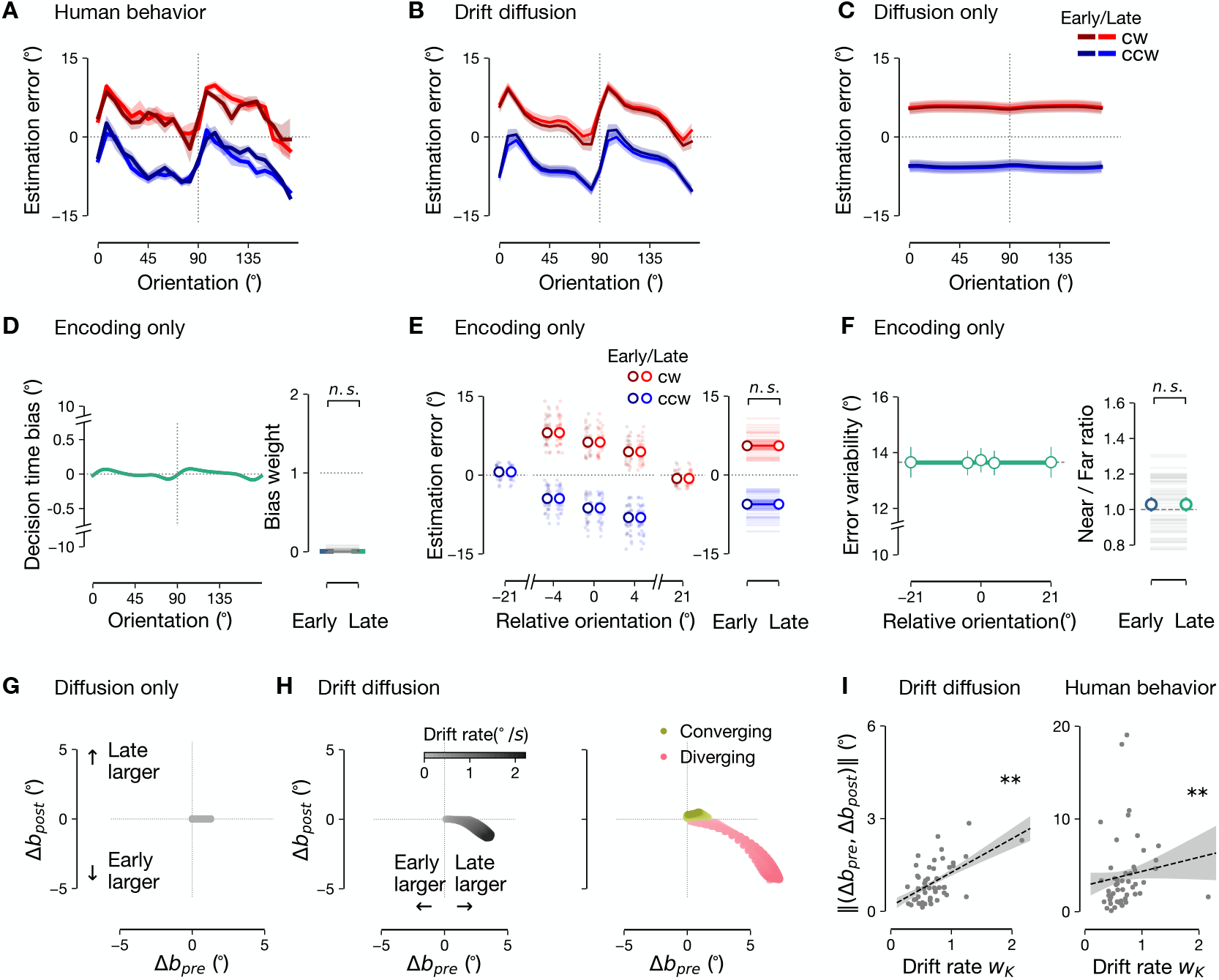
Analysis of biases in behavior and phenomenological models, Related to Figures 3 and 4. **A-C**, Stimulus-conditioned decision-consistent biases in behavior and phenomenological models. **A**, Behavioral estimation errors, conditioned on each choice and stimulus, in the early (darker) and late (brighter) decision conditions. Colored shades denote mean ± s.e.m.s across trials. **B,C**, Model predictions of estimation errors with the best-fitting parameters for each participant, in the drift-diffusion model (**B**) and the diffusion-only model (**C**). **D-F**, Model predictions of stimulus-specific bias (**D**), decision-consistent bias (**E**), and near-reference variability **(F)** under encoding-only (without drift nor diffusion) dynamics. Format, same as in **Figures 3D-F. G-I**, Drift modulation of pre- and post-decision biases. **G,H**, *Ex-ante* simulation of the decision-timing-dependent changes in the pre-decision 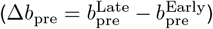 and post-decision 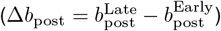 biases under the diffusion-only **(G)** and the drift-diffusion (**H**) dynamics. In **G** and the left panel of **H**, the data were merged across all stimulus orientations. In the right panel of **H**, the data are shown separately for the orientations with diverging and converging drift. **I**, Relationship between the estimated drift rate parameter (*w*_*K*_) and the distance of (Δ*b*_pre_, Δ*b*_post_) from the origin, in the drift-diffusion model *ex-post* simulation (left) and in human behavior (right). Spearman’s *ρ* = 0.422, *p* = 0.0023 (left), *ρ* = 0.385, *p* = 0.0057 (right). * * \**p <* 0.001, * \**p <* 0.01, ns *p* > 0.05.

**Figure S4.**
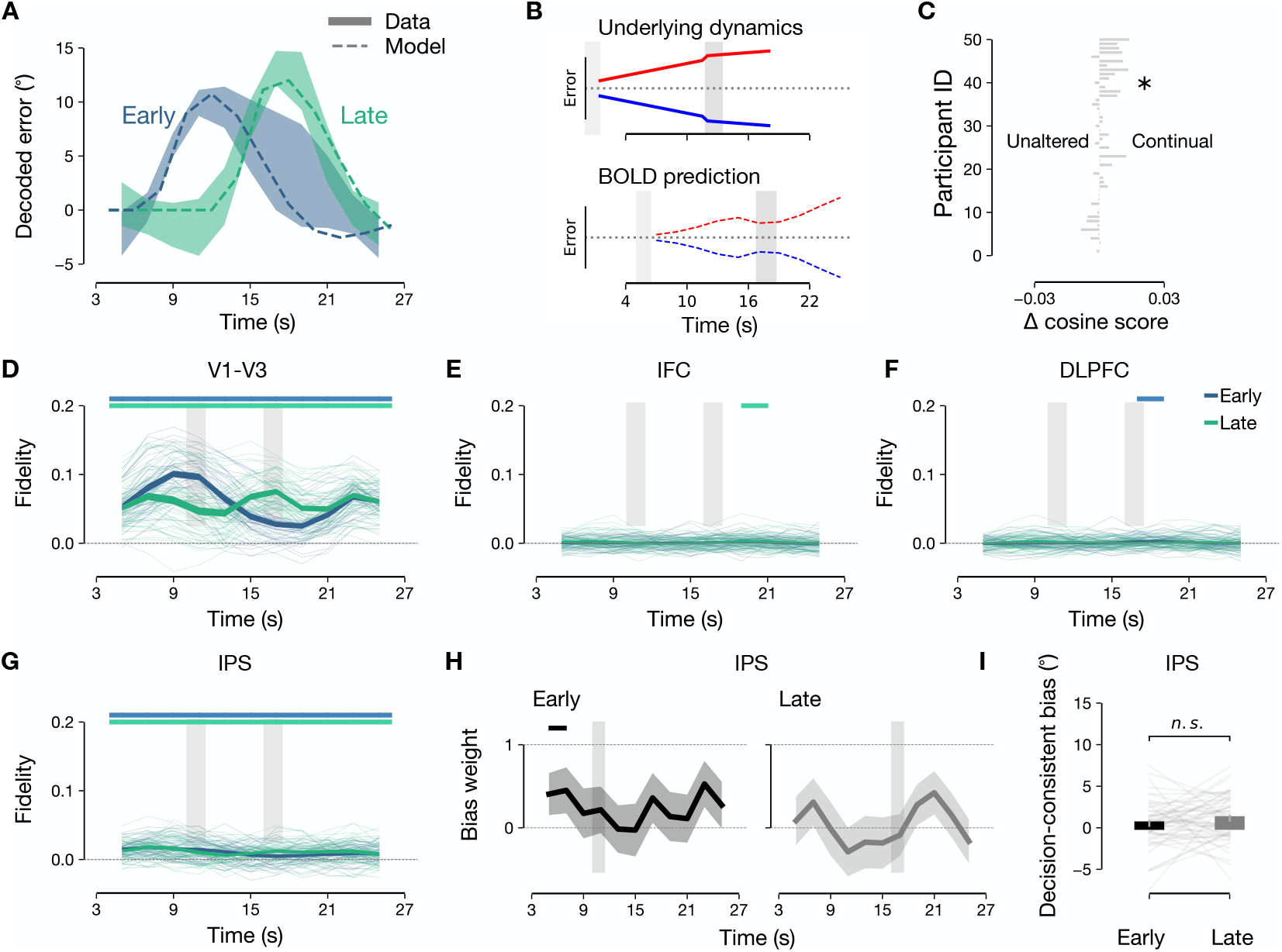
Analyses of BOLD responses, Related to Figure 5. **A**, Decoded errors (V1-V3) for far-reference trials used to estimate the beta weights for stimulus (*β*_*θ*_) and reference (*β*_*ρ*_) using an event-related regression analysis on the circular space (see **STAR Methods**). Shades denote bootstrapped data. **B**, Convolution in the polar space, combined with visual drives, converts the underlying memory dynamics (top) into a BOLD dynamics (bottom) prediction. Note there is a transient dip in the predicted dynamics due to the visual drive from the reference, which is not present in the underlying memory dynamics. **C**, Parameter-free comparison between the cosine scores (evaluated against the BOLD decoding trajectory) of the continual updating of the drift-diffusion model and the encoding-only, unaltered memory model. Participants are sorted as in **Figure 5f**. Paired *t*-test, *t*(49) = 2.483, *p* = 0.0165. **D-G**, Decoding fidelity [S4] of the stimulus orientation for each time point in V1-V3 (**D**), IFC (**E**), DLPFC (**F**), and IPS (**G**). **H,I**, Same as **Figures 5B,D**, but using the decoded memory states from IPS. In **D-H**, horizontal bars above the panels denote the time points where decoding fidelity is significantly different from zero (permutation test, Bonferroni-corrected for number of time points). * *p*< 0.05, ns *p*> 0.05.

**Figure S5.**
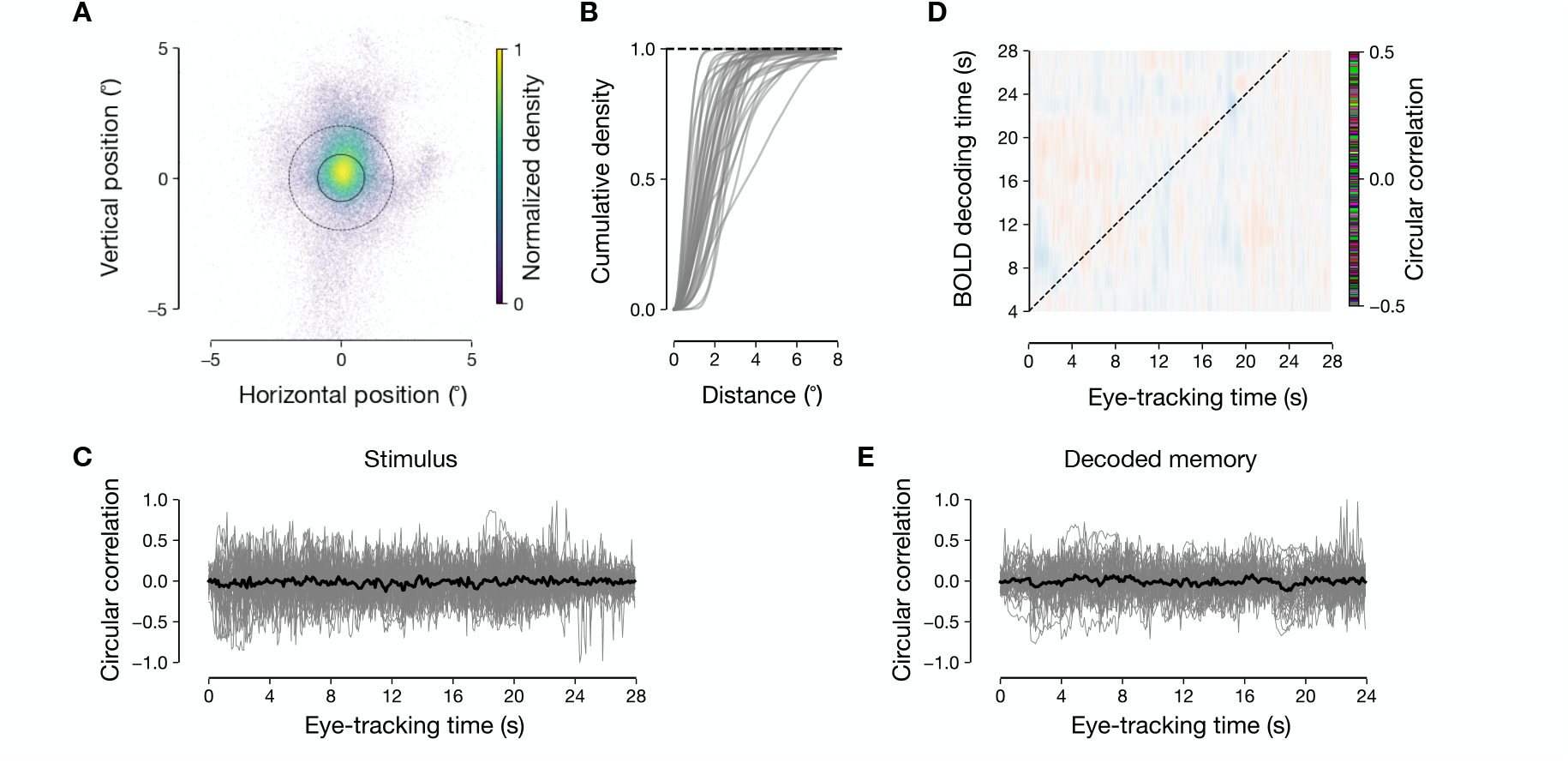
Eye-tracking data analysis, Related to Figure 5. **A**, Eye positions across participants during the experiment sampled at 500Hz. Solid inner ring denotes the fixation ring size, and the dashed outer ring denotes the inner aperture of the oriented grating stimulus. **B**, Cumulative density of eye position distribution as a function of distance from the center. Each gray line denotes a participant. **C**, Circular correlations between stimulus orientation and angular eye positions for each time point within a trial (gray line, individuals; thick black line, population average; *p* > 0.05 for all time points, FDR-corrected one-sample *t*-test against zero). **D**, Circular correlations between BOLD decoded orientations (using inverted encoding analysis) and angular eye positions for combinations of time points within a trial (*p*> 0.05 for all cells, FDR-corrected one-sample *t*-test against zero). **E**, Same as **C**, but circular correlations between BOLD decoded orientation and angular eye positions for their corresponding time points considering hemodynamic delay of 4s, denoted as dashed diagonal line in **D** (*p*> 0.05 across all time points, FDR-corrected one-sample *t*-test against zero).

**Figure S6.**
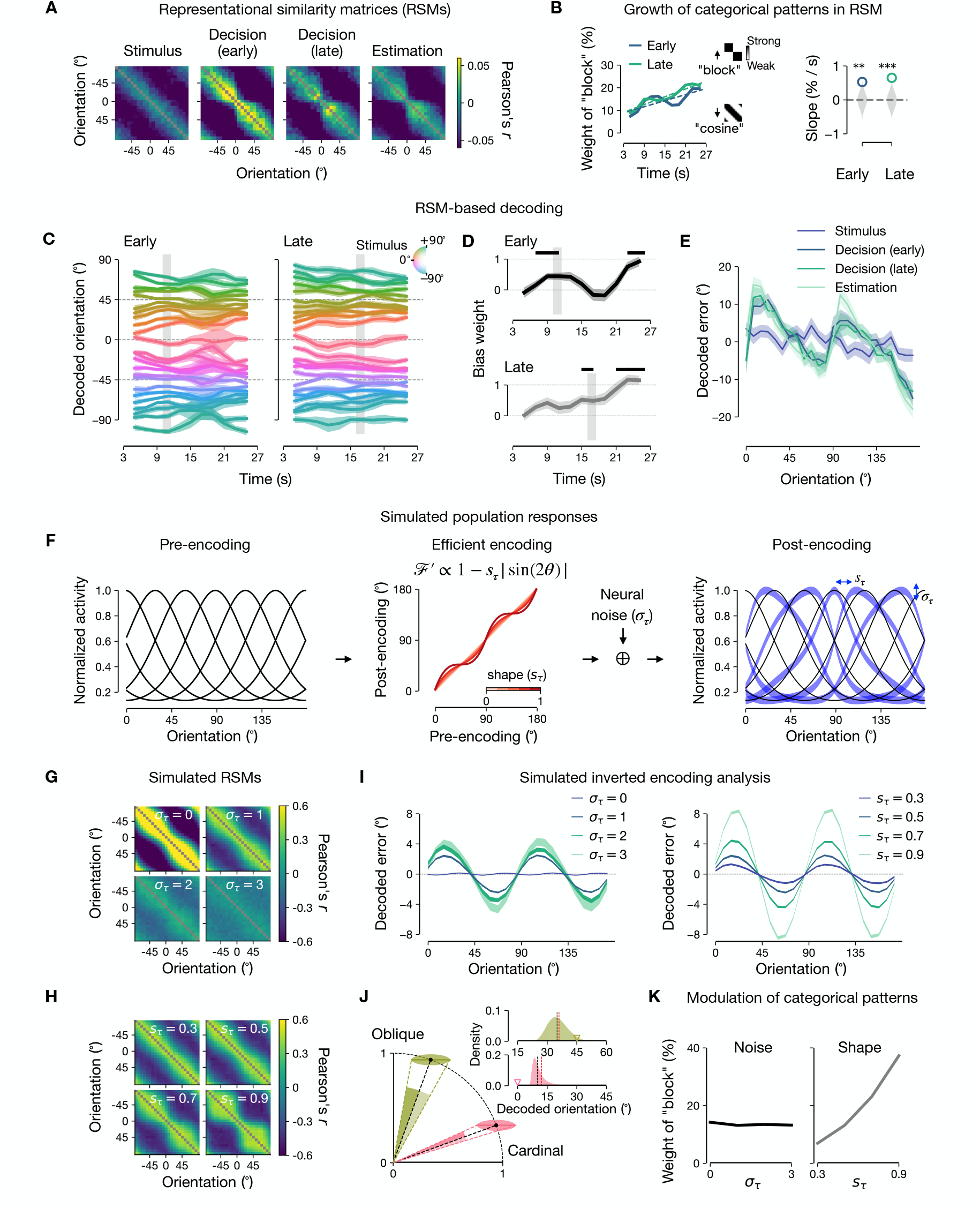
Representational similarity analysis and decoding of simulated population responses, Related to Figure 5. **A**, Representational similarity matrices (RSM) based on Pearson correlation coefficients at four landmark task time points in temporal order, using the same voxel population used for the inverted encoding analysis. **B**, Left, Progression of relative weights (in percent) of the hypothesized “block” matrix (upper inset) compared to “cosine” circulant matrix (lower inset) patterns in explaining the shapes of time point-wise RSMs, measured by determining the convex combination weights between the two patterns (see **STAR Methods**). Right, linear slope (in %*/s*) estimates of the growth of relative weights over time. Permutation test, *p* = 0.0031 (Early), *p <* 10^−4^ (Late). **C-E**, RSM-based decoding analyses, where the average memory state corresponding to each stimulus is estimated via circular averaging of each row of RSMs for each time point. **C,D**, Gray vertical bars demarcate the time of discrimination in the early and late discrimination trials, taking into account the canonical hemodynamic delay of 4s. **C**, Temporal trajectories of RSM-decoded memory states, pooled across participants: colors, stimulus orientations, shades, ± s.e.m.s based on bootstrapping. **D**, Bias weight, measured as the regression coefficient of the RSM-decoded errors against each participant’s stimulus-specific bias function, for early (upper) and late (lower) conditions: shades, ± s.e.m.s across participants. Black horizontal bars at the top demarcate the time points over which the mean of the bias weight significantly differed from zero (*p <* 0.05, one-sample *t*-test, Bonferroni-corrected for the number of time points). **E**, Stimulus-specific errors of the decoded orientations for four landmark time points in **A**. ± bootstrap s.e.m. **F**, Simulated population responses based on an influential implementation of efficient encoding [S5]. Symmetric tuning distribution (left panel) is transformed by the efficient encoding function [S5–7] whose levels of heterogeneity depend on the shape parameter *s*_*τ*_ (middle panel). The post-encoding population responses after adding Gaussian noise (*σ*_*τ*_) [S8–10] were used as simulated voxel responses. **G,H**, Representational similarity matrices of the simulated population with different parameters (for **G**, different choice of noise parameters while *s*_*τ*_ is fixed at 0.5; for **H**, different choice of shape parameters while *σ*_*τ*_ is fixed at 1). **I**, Errors of the decoded orientations via the inverted encoding analysis under different additive noise (*σ*_*τ*_), and different shape (*s*_*τ*_) based on the same cross-validation procedure we used for the BOLD data. In the absence of noise (left, *σ*_*τ*_ = 0), decoding error remains unbiased even with heterogeneously tuned population responses, as the decoder already accounts for this heterogeneity. However, in the presence of noise, decoding error becomes increasingly biased with greater noise levels (left) or tuning heterogeneity (right). **J**, Schematic of how noise and shape jointly contributes to the decoding of biases. Colored ellipses denote the projections of the estimated channel responses for a stimulus near cardinal (pink) and near oblique (green) orientations. The inverse of the tuning similarity matrix (ℐ ^⊤^ℐ) ^−1^ (where ℐ denotes the tuning matrix of the shape [*N* _units_ × *N*_stimuli_] from **F**) is used to estimate channel responses and endows horizontally amplified noise to the estimated channel responses when projected onto the unit circle (see **Method S2.1**). The orientation readouts from the projections (corresponding inset panels), as done in the inverted encoding analysis, lead to a pattern of *cardinal repulsion* with an attractive bias towards the nearby oblique orientations (green inverted triangle) and a repulsive bias from the nearby cardinal orientations (pink inverted triangle): black vertical lines, true stimulus, red vertical lines, average of the simulated decoded stimuli from inverted encoding analysis. **K**, Weights of the “block” pattern in the two different simulation conditions (varying noise levels via *σ*_*τ*_ and varying shapes via *s*_*τ*_). ^* * *^*p <* 0.001, ^* *^*p<* 0.01.

**Figure S7.**
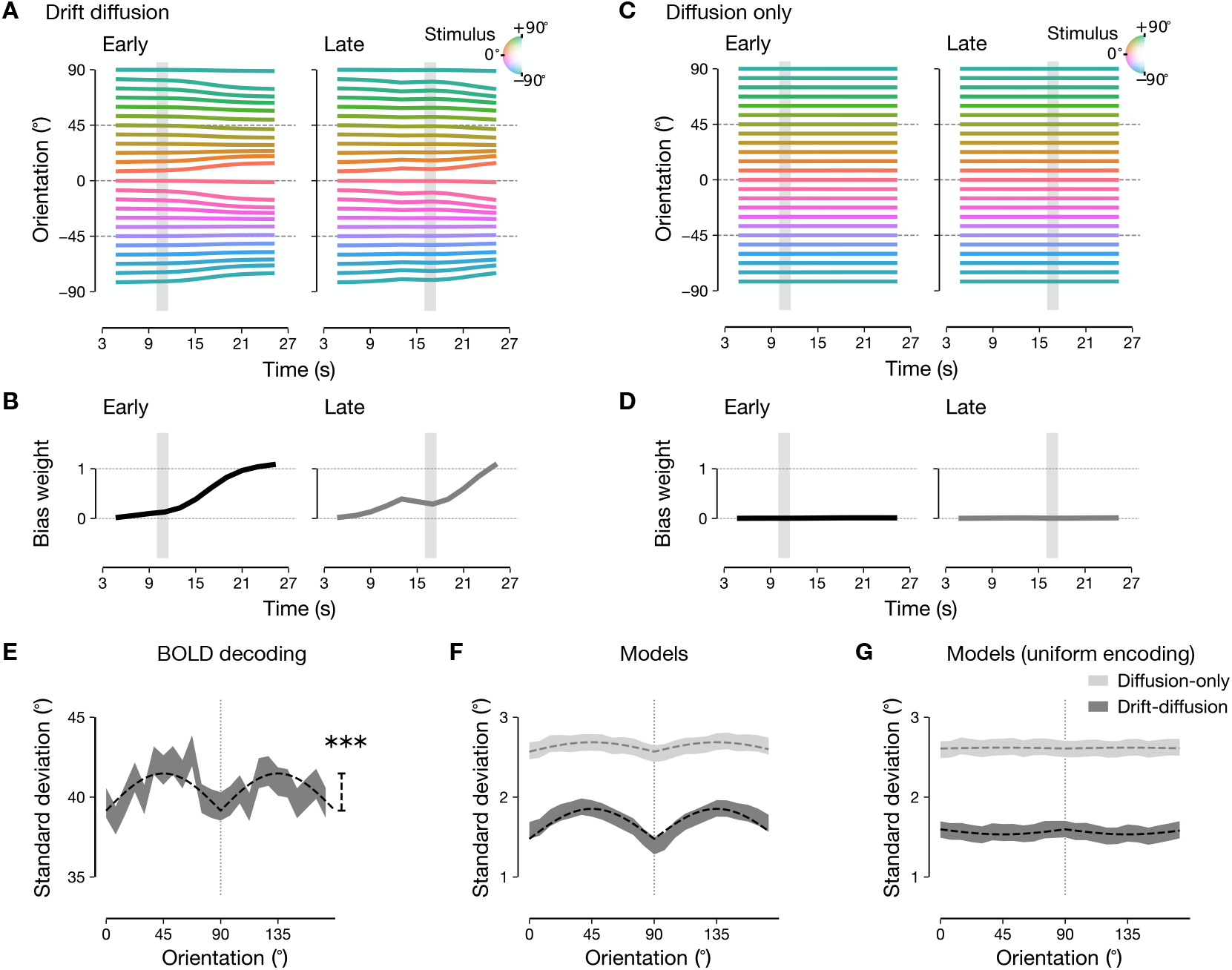
Phenomenological model prediction of BOLD responses, Related to Figure 5. **A**, Evolution of memory states in early (left) and late (right) DM conditions in the drift-diffusion model, conditioned on each stimulus orientation (color). Shown are the *ex-post* model simulations based on the best-fit parameters of the drift-diffusion model for individual participants and the hemodynamic convolution in the polar space (see **STAR Methods**). **B**, Evolution of the stimulus-specific bias weight, with a value of 1 corresponding to the amplitude of the behavioral bias predictions. **C-D**, Same as **A-B**, but with the *ex-post* model simulations based on the best-fit parameters of the diffusion-only model. **E-G**, Decoding of BOLD responses around encoding time. **E**, Across-trial standard deviation of BOLD decoding of memory states around the stimulus presentation (“encoding time”, TRs falling within 4s to 8s after the stimulus onset), conditioned on each stimulus orientation. Data were pooled across participants, ± bootstrap s.e.m. Dashed line denotes the fitted rectified sine curve, *α*_*E*_ |sin 2*θ*| + *β*_*E*_, following Girschick *et al*. [S11]. The vertical line on the right denotes the best-fit amplitude, *α*_*E*_ = 2.325°. Permutation test, *p* = 0.0002. **F**, Across-trial standard deviations of the memory states predicted by drift-diffusion and diffusion-only models, convolved with the hemodynamic response function and probed at the same time window as fMRI data. ± bootstrap s.e.m. **G**, Same as **F**, but with the encoding shape parameter (*s*) set to 1, corresponding to the uniform stimulus-to-sensory mapping (see **STAR Methods**). ± bootstrap s.e.m. ^* * *^*p<* 0.001.

**Figure S8.**
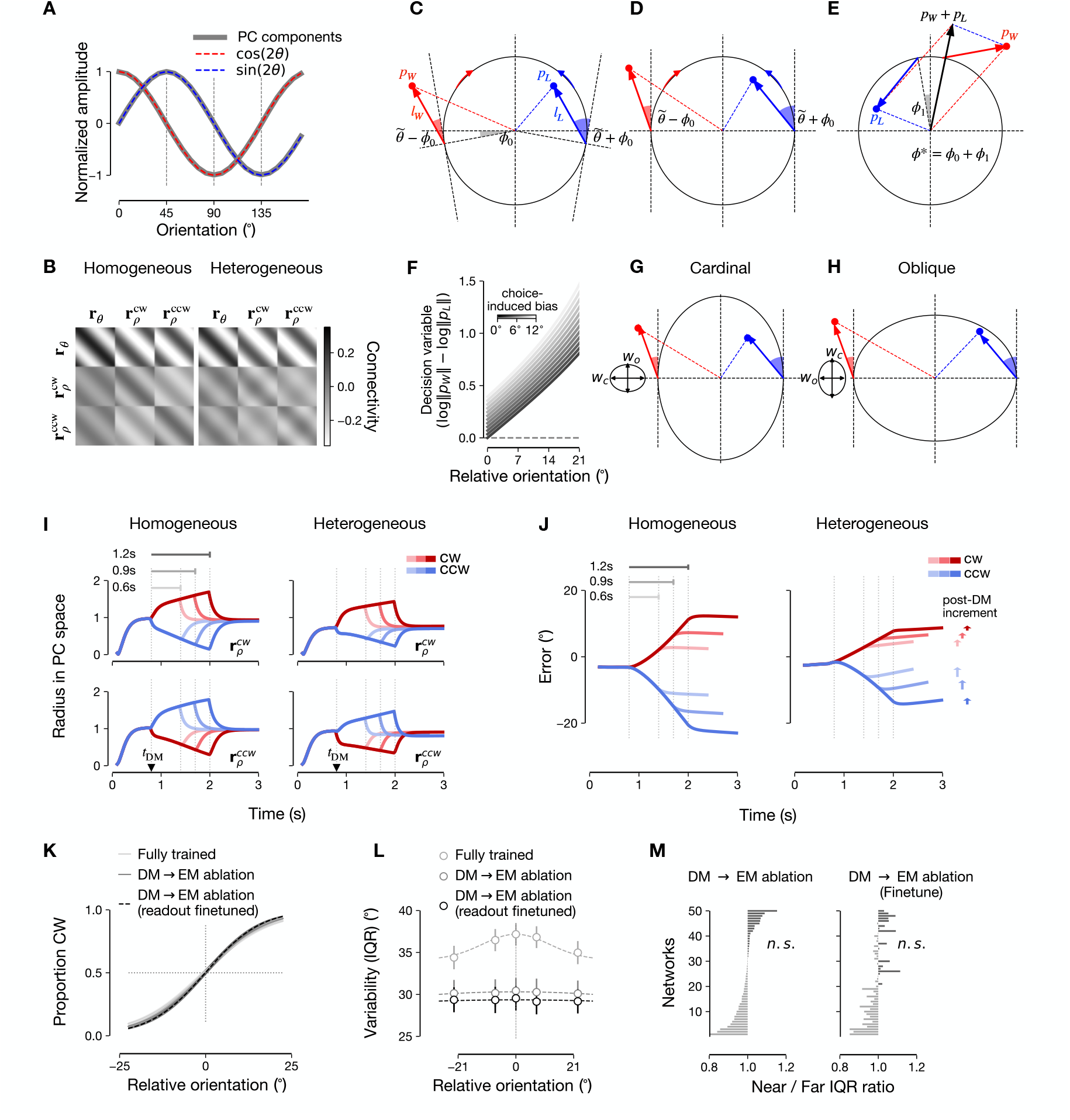
Analysis of choice-induced bias in trained RNNs, Related to Figures 7 and 8. **A**, Similarity between the columns of the principal component projection matrix **V**_𝒟_ (solid lines) and the cosine and sine curves (dashed lines). **B**, Rank-3 approximations of the blocks of the trained ***J*** (see **Method S4.1**). **C-E**, Linear construction of choice-induced bias on a unit circle. **C**, Reference input direction 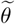 forms angles 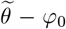 with the tangent of the winning population vector (red) and 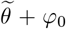 with that of the losing population vector (blue), where φ_0_ denotes ***r*** _*θ*_ *→* ***r*** _*ρ*_ displacement angle beyond *π /*2. In general, the reference responsivity (changes in the populations due to the reference input) can differ between winning (*l*_*W*_) and losing (*l*_*L*_) populations. **D**, Rotation of the vectors in **C** by ± *π /*2 ± *φ*_0_, yielding an equivalent setup as **C**. Note that both winning and losing populations start from the x-axis, but the responding angles differ. **E**, Rotation of vectors in **D** by ∓ *π/*2 *φ*_1_, where *φ*_1_ denotes ***r*** _*θ*_ → ***r*** _*ρ*_ displacement angle (**Figure 7G**). The addition of the winning population vector *p*_*W*_ and losing population vector *p*_*L*_ yields the biased stimulus representation updating direction *p*_*W*_ + *p*_*L*_. We show that the bias (the tangent component of *p*_*W*_ + *p*_*L*_) is decision-consistent when *φ*^*^ = *φ*_0_ + *φ*_1_ > 0 and *l*_*W*_ ≥ *l*_*L*_ (see **Method S4.2**). **F**, An advantage of having a choice-induced bias. Decision variable log ‖ *p*_*W*_ ‖*/* ‖ *p*_*L*_‖, the difference between the norms of *p*_*W*_ and *p*_*L*_, following the choice-induced bias, becomes larger compared to that without a choice-induced bias, leading to more correct decisions possibly under noise. **G-H**, The impact of warped rotations in heterogeneous RNNs, using the setup corresponding to **D. G**, Around the cardinal orientations, due to the geometry of the warped rotation imposed by ***J***, the elongation along the direction normal to the ellipse (*w*_*c*_) is larger than the elongation along the tangential direction (*w*_*o*_). **H**, Around the oblique orientations, elongation along the tangential direction (*w*_*c*_) is larger than the normal direction (*w*_*o*_). **I-J**, Pulse-like impact of DM on memory states in trained RNNs. **I**, Transient choice formation induced by reference input 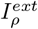 in the reference-receiving population (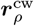 and 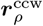). Durations of the discrimination epoch were varied as 0.6s, 0.9s, and 1.2s and the radii of the reference population responses in the PC space were measured. For homogeneous RNNs (left panel), the reference input during the discrimination epoch induces the expansion of the to-be-selected (winner) subpopulation, and the contraction of the not-to-be-selected (loser) subpopulation, with an almost symmetric amount of radius deviations. After the reference input is gone, the expansion and contraction quickly relaxes back to their original levels. For heterogeneous RNNs (right panel), similar dynamics were observed, but with an asymmetric amount of radius deviations. Here, we used the stimulus near (± 15 °) to the cardinal orientation (*θ* = 90°). **J**, Impact of the choice-induced bias in memory states for estimation. Even though the choice-related changes in the reference-receiving populations were short-lived, the immediately updated memory representation in ***r***_*ω*_ persists after the disappearance of the reference. For the homogeneous RNNs (left panel), the amount of decision-consistent bias is modulated by the duration of reference inputs. Yet, without drift dynamics, there is a minimal post-DM increment, *i*.*e*., *b*_*post*_, subtracted by the transient amount from choice-induced bias. For the heterogeneous RNNs (right), the amount of decision-consistent bias is modulated not only by the duration of reference inputs but also by drift. **K**, Discrimination psychometric curves of fully trained RNNs (light gray curve), RNNs with the feedback connection, **r** _*ρ*_ → r _*θ*_, ablated (darker gray curve), and the same ablated networks whose readout weights were further finetuned with 100 more iterations (black curve). **L,M**, No increase of near-reference variability in feedback-ablated networks and their finetuned counterparts. ns *p* > 0.05.

**Table S1.** Phenomenological model parameters and their constrained ranges, Related to STAR Methods.

| Symbol | Parameter | Optimization bounds |
| --- | --- | --- |
| $w_K$ | Drift rate for memory | $[0, 15] (^{\circ}/s)$ |
| $w_D$ | Diffusion rate for memory | $[0, 15] (^{\circ}/\sqrt{s})$ |
| $w_{\alpha}$ | Choice-induced bias | $[-15, 15] (^{\circ})$ |
| $w_{\rho}$ | Reference bias weight | $[0, 15]$ |
| $w_E$ | Encoding variability | $[0, 15] (^{\circ})$ |
| $w_P$ | Production variability | $[0, 15] (^{\circ})$ |
| $w_{\lambda}$ | Decision-making lapse rate | $[0, 0.15]$ |
| $s$ | Encoding function shape parameter | $[0, 1]$ |

## Method S1 Analysis of behavior data

### Method S1.1. Derivation of decision-consistent biases, Related to Figure 4 and STAR Methods

Here, we analytically expressed the pre- and post-decision components of the decision-consistent bias, Δ*b*_pre_ and Δ*b*_post_. For *b*_pre_, we used the Bayes rule to derive the conditional probability of the memory state *m*_*t*_ at pre-decision time *t* ≤ *t*_dm_, given the choice *ĉ*_*λ*_ with the lapse rate *λ*, along with the stimulus *θ* and the reference *ρ*.

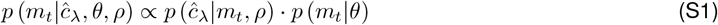

The first term on the right-hand side is

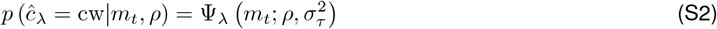

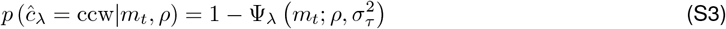

where Ψ_*λ*_ is the Gaussian psychometric function and *σ*_*τ*_ is the amount of the noise level accumulated from time *t* to the decision time *t*_dm_. The second term is 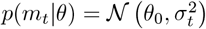. For each choice, the conditional means of Equation (S1) are given as

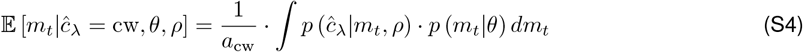

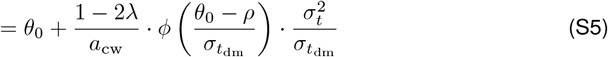

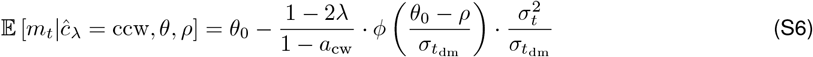

where 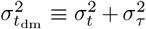 and the normalization constant 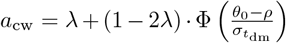. Thus, setting *t* = *t*_dm_, *b*_pre_ is expressed as

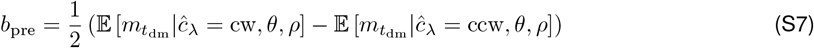

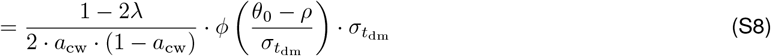

Note that we obtained 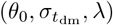 by fitting the psychometric function Ψ_*λ*_ to the decision-making data. Thus, only the decision-making behavior is required to estimate *b*_pre_.

### Method S1.2. Near-reference bias, Related to Figure 8

We estimated near-reference bias (increased bias of marginal error distribution when the stimuli are near references), characterized as indicative of choice-induced bias [S2], We expressed the near-reference bias as a weight of choice *w*_*α*_, while regressing out the impacts of reference attraction *w*_*ρ*_. We extended the equations in Method S1.1 to the memory state right after DM, denoted as 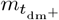, which is updated by *α* as

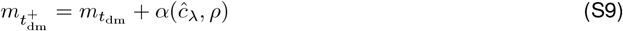

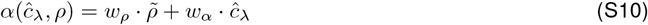

where 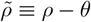. We have the conditional means at 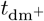 as follows.

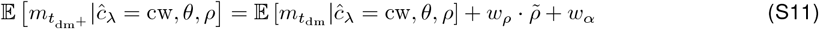

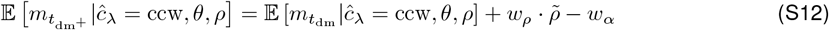

Next, we replace the representation for the marginal mean with the equations (S11)-(S12) as follows.

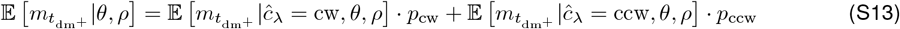

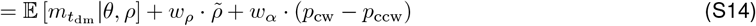

where *p*_cw_ ≡ *p* (*ĉ*_*λ*_ = cw|*θ, ρ*) and *p*_ccw_ = 1 − *p*_cw_. Thus, we can characterize the reference attraction 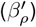 and the choice-induced bias 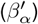 as the coefficients of the following linear regression.

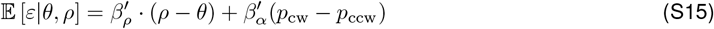

## Method S2 Analysis of BOLD data

### Method S2.1. Robustness of bias decoding from BOLD signals, Related to Figures 5 and S6

To test whether the stimulus-specific bias is decodable using the inverted encoding analysis, we simulated a neural population activity following an influential efficient coding model by [S5] (**Figure S6F**).

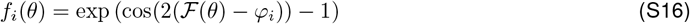

with the activity of the unit *i* with the preferred orientation *φ*_*i*_ and the encoding function F of the stimulus *θ*. Following the assumptions made in [S6] to construct the encoding function from natural statistics prior, we used a simple encoding function ℱ with the functional form [S12] with a parameter 0 ≤ *s*_*τ*_ ≤ 1,

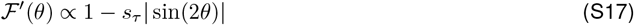

where lower *s*_*τ*_ corresponds to the “flatter” slope of the encoding function ℱ. The slope of the encoding function captures the amount of neural coding resource [S13].

To simulate the modulation of the population activity, we assumed two scenarios: noise and tuning hetero-geneity (“shape” hereafter). For noise modulation, we added the Gaussian noise of *σ*_*τ*_ to *f*_*i*_(*θ*), following frequently made assumptions in fMRI literature [S8–10]. Using the equivalent decoding procedure with the inverted encoding analysis used for human BOLD decoding, we found that sinusoidal bias patterns were observed, increasing with the noise level *σ*_*τ*_ and shape *s*_*τ*_ (**Figure S6I**). For simulation, we used *σ*_*τ*_ ∈ {0, 1, 2, 4} for *s*_*τ*_ = 0.5, and *s*_*τ*_ ∈ {0.3, 0.5, 0.7, 0.9} for *σ*_*τ*_ = 1. We used 1358 units (median number of used voxels in humans), with the same number of trials as human experiment, and we repeated 50 times, to compute the standard errors, matching the number of participants.

We investigated how such biases could arise from both sources, noise or shape, assuming an idealized case. Consider, without loss of generality, a tuning matrix of ℐ is a biased representation that depends on the tuning shape *s*_*τ*_. Concretely, let T to be [*N*_unit_, *N*_channel_] computed from the training set, assuming the same number of observations per channel in the training set. Here, as we use equally spaced stimuli, *N*_channel_ = *N*_stimuli_. A voxel-wise encoding model for a specific stimulus is given as **b** = ℐ**c**, thus the inversion is given as ***ĉ*** = (ℐ^T^ℐ) ^—1^ ℐ^T^**b**.

Now, for a test-set trial of stimulus *k*, assume an independent, Gaussian additive noise to the population response vector, **b**_*k*_ = t_*k*_ + ***ε***_*k*_. Then the channel response ***ĉ*** is given as ***ĉ***_*k*_ = **e**_*k*_ + (ℐ^T^ℐ) ^—1^ ℐ^T^***ε***. Thus,

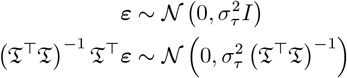

With sinusoids **s** and cosinusoids **n**, we have the projections of the channel response to get the stimulus decoding 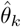,

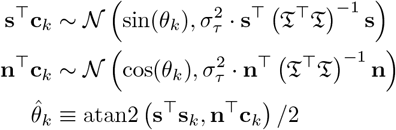

Note that in the case of no noise, 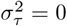, the decoding 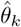 is the same as *θ*_*k*_, showing no bias. In the presence of noise, how the 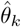 is affected by noise depends on the interaction between sinusoids and ℐ. In the choice of ℐ respecting [S5], **n**^T^ (ℐ^T^ℐ) ^− 1^ **n** dominates **s**^T^ (ℐ^T^ℐ) ^− 1^ **s**, generating an ellipsoidal noise distribution on a decoding circle, on which each estimated channel is projected (**Figure S6J**). Thus, such biased noise distribution leads to a sinusoidal bias in decoding.

## Method S3 Phenomenological model

### Method S3.1. Numerical procedure for model fitting, Related to STAR Methods

To fit the drift-diffusion model represented as a Fokker-Planck equation in Equation 22, we used the finite difference method with *N*_disc_ = 96 discretization points with Δ*m* = *π/N*_disc_,

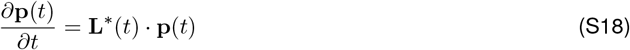

where **p**(*t*) is discretized *p*(*m, t*). The transition matrix **L**^*^(*t*) denotes the following,

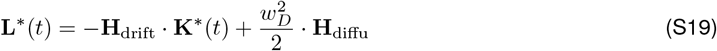

where **K**^*^(*t*) is the diagonal matrix with the diagonal elements corresponding to discretized *K*^*^(*m, t*), and **H**_drift_ and **H**_diffu_ denote the (*N*_disc_ × *N*_disc_) matrices with the elements equal to zero except for the following,

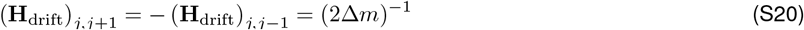

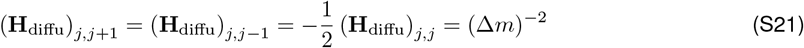

with the *N*_disc_-periodic subscripts for the wrapping-around orientations. For convenience, we define the time-independent transition matrix **L** for every time point except *t*_dm_.

To compute the full density of the memory process, using the efficient sensory encoding constrain, we first computed post-encoding distribution **p**_0_, which is a vector representing the densities *p*(*m*_0_|*θ*) for the given *θ*, evaluated at the discretization points of *m*. Next, to evaluate the density of working memory at *t*_DM_, we computed **p**_DM_, a vector representing the densities 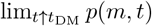, as follows.

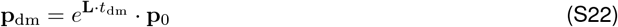

To compute the conditional distributions after decision-making, we defined a set of mask functions,

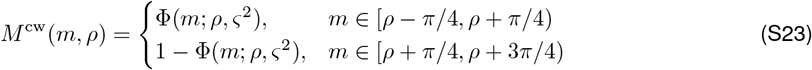

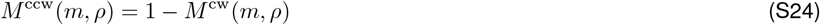

where Φ denotes the Gaussian cumulative density function, and *ς* = 0.2 captures the amount of the noise associated with the reference, which results in the smoothness of decision conditioning. Next, we applied the masks and choice-induced bias with strength *w*_*β*_ to **p**_dm_,

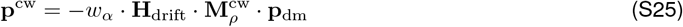

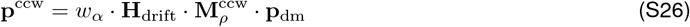

with 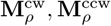 denote the diagonal matrices with the elements corresponding to *M* ^cw^(*m, ρ*),*M* ^ccw^(*m, ρ*) evaluated at the discretization points of *m*. Additionally, we added the decision lapse rate parameter *w*_*λ*_, resulting in a modified set of conditional distributions.

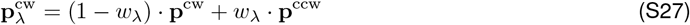

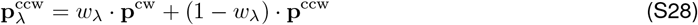

Finally, we propagated these conditional distributions to *t*_EM_ and convolved them with the production error associated with the estimation report, obtaining the joint density of discrimination and estimation,

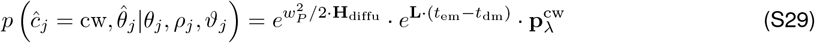

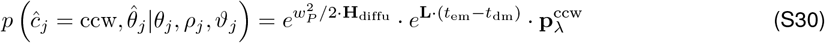

where *w*_*P*_ is the width of the production error distribution, and *θ*_*j*_, *ρ*_*j*_, *θ*_*j*_ denote each trial *j*’s stimulus and reference orientations, and decision-making condition, respectively.

## Method S4 Recurrent neural network model

### Method S4.1. Low-dimensional description of RNN state space, Related to Figures 7-8 and S8

To describe the dynamics underlying the choice-induced bias, we tightly approximated the columns of the principal component projection matrix **V**_𝒟_ with sine and cosine functions **Figure S8A**, forming the basis matrix **V** for *j* = 1, ···, 24.

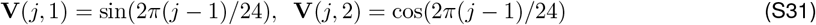

For a low-dimensional description of the trained connectivity **J**, we reduced the dimension for ***r***_*θ*_ from 48 into 24 by averaging the block matrices corresponding to 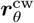 and 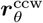. This resulted in nine (24 × 24) blocks, representing the projections between 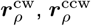, and ***r***_*θ*_. The RNN dynamics without noise during the reference presentation are approximated as follows.

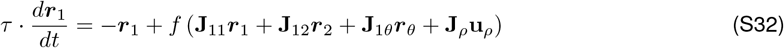

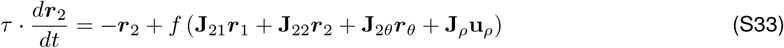

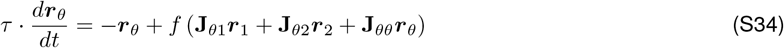

where ***r***_1_, ***r***_2_ denote 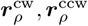, respectively, and **J**_11_, ··· **J**_*θθ*_ are the corresponding block matrices.

To describe the dynamics of the low-dimensional vectors ***s***_*j*_ ≡ **V**^T^***r***_*j*_, we approximated each block as the following rank-3, weighted circulant matrix form by minimizing the Frobenius norm ‖ **J**_block_ − **Ĵ** _block_‖ _*F*_ (**Figure S8B**).

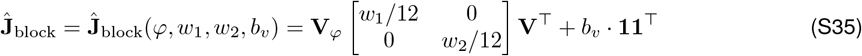

where **V** _*φ*_ is the basis matrix with the phase(*φ*)-shifted sine and cosine functions.

With the rank-3 form, we exploited the following relationship.

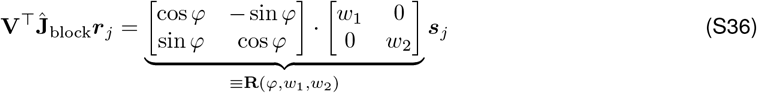

where **R** = **R**(*φ, w*_1_, *w*_2_) describes the warped rotation such that ***s***_*j*_ is warped by *w*_1_ and *w*_2_ along each PC and then rotated by *φ* in a canonical direction. As PC1 and PC2 axes correspond to cardinal and oblique stimuli axes in ***r***_*ρ*_, we semantically denote *w*_*c*_ = *w*_1_ and *w*_*o*_ = *w*_2_ in the main text.

With the warped rotation revealed by **Ĵ**, we further approximated the low-dimensional dynamics. Using a linear function 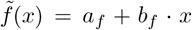 instead of *f* in equations (S32)-(S34), we obtained the following reduced dynamics in ***s***_1_, ***s***_2_, ***s***_*θ*_, each corresponding to two-dimensional projections of ***r***_1_, ***r***_2_, ***r***_*θ*_.

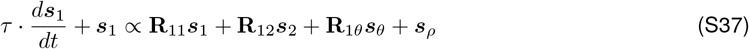

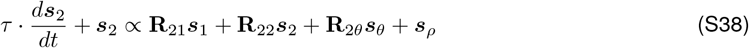

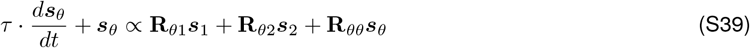

where ***s***_*ρ*_ ∝ [sin *ρ*, cos *ρ*]^T^ denotes the reference input direction. As such, this formulation justifies our interpretation of the recurrent connections (**Figures 7A,B**) as rotation-addition.

### Method S4.2. Linear analysis of choice-induced bias in RNNs, Related to Figures 7-8 and S8

We next explain how choice-induced bias arises in the reduced and linearized dynamics in equations (S37)-(S39). For simplicity, we use the two-dimensional vectors *p*_*W*_ and *p*_*L*_ to represent the activities of winning and losing reference-receiving populations. The log ratio of their lengths, DV = log ‖ *p*_*W*_ ‖*/* ‖ *p*_*L*_ ‖, serves as a decision variable, characterizing the relative amplitudes of the expansion and contraction in the RNN vectors. We first assumed equal changes in reference input (*i*.*e*., reference responsivity), denoted by *l*_*W*_ and *l*_*L*_, while they may differ due to RNN nonlinearity.

In homogeneous RNNs with *w*_*c*_ = *w*_*o*_, we found that 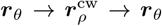 can be represented as rotations by −*π/*2 − *φ*_0_ and +*π/*2 + *φ*_1_ in sequence for small *φ*_0_ and *φ*_1_ (signs are flipped for CCW; **Figures S8C-E**). With relative orientation 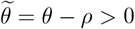 and 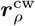 as the winning population, an external reference input forms an angle 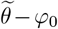 with the tangent in 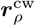, and 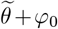 in 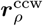 at the onset of the DM task (**Figures S8C,D**). After rotation back *r* _*ρ*_ → *r*_*θ*_, the contributions from 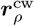 and 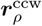 are summed as *p*_*W*_ + *p*_*L*_ (**Figure S8E**). Its tangential component is

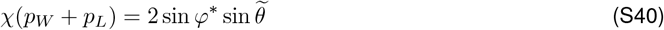

where *φ*^*^ = *φ*_0_ + *φ*_1_ is the total rotation displacement angle in ***r***_*θ*_ → ***r***_*ρ*_ → ***r***_*θ*_ rotations. For 0 <*φ*^*^ <*π* (as in our trained RNNs), this results in a sinusoidal, *repulsive* form of choice-induced bias as a function of 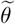.

For the heterogeneous RNNs, the warping weights, *w*_*c*_ and *w*_*o*_ differ, causing anisotropic transformations between ***r***_*ρ*_ and ***r***_*θ*_. Specifically, *w*_*c*_ >*w*_*o*_ for both **R**_*θ*1_, **R**_*θ*2_ (whose warping weights are fit to be near-identical), resulting in larger expansion in the normal direction than in the tangential direction around cardinal orientations (**Figure S8G**; opposite around the oblique orientations in **Figure S8H**). Equation (S40) is modified as

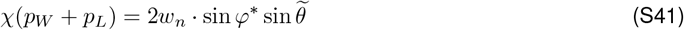

where *w*_*n*_ = *w*_*c*_ for cardinal stimuli and *w*_*n*_ = *w*_*o*_ for oblique stimuli, leading to a larger bias around cardinal orientations than oblique orientations (**Figures 8F,G**).

Finally, when *l*_*W*_ and *l*_*L*_ are different, Equation (S41) generalizes to

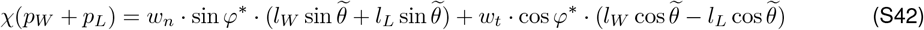

where *w*_*n*_ and *w*_*t*_ are the warping weights along the normal and tangential directions of the ellipse, respectively. Under 0 <*φ*^*^ <*π* and *l*_*L*_ ≤ *l*_*W*_, the repulsive bias emerges.

## References

1. Ma, W.J., and Jazayeri, M. (2014). Neural coding of uncertainty and probability. Annu. Rev. Neurosci. 37, 205–220. 10.1146/annurev-neuro-071013-014017.

2. Jazayeri, M., and Movshon, J.A. (2007). A new perceptual illusion reveals mechanisms of sensory decoding. Nature 446, 912–915. 10.1038/nature05739.

3. Gold, J.I., and Shadlen, M.N. (2007). The Neural Basis of Decision Making. Annu. Rev. Neurosci. 30, 535–574. 10.1146/annurev.neuro.29.051605.113038.

4. Mante, V., Sussillo, D., Shenoy, K. V, and Newsome, W.T. (2013). Context-dependent computation by recurrent dynamics in prefrontal cortex. Nature 503, 78–84. 10.1038/nature12742.

5. Bogacz, R., Brown, E., Moehlis, J., Holmes, P., and Cohen, J.D. (2006). The physics of optimal decision making: A formal analysis of models of performance in two-alternative forced-choice tasks. Psychol. Rev. 113, 700–765. 10.1037/0033-295X.113.4.700.

6. Romo, R., Brody, C.D., Hernández, A., and Lemus, L. (1999). Neuronal correlates of parametric working memory in the prefrontal cortex. Nature 399, 470–473. 10.1038/20939.

7. Machens, C.K., Romo, R., and Brody, C.D. (2005). Flexible Control of Mutual Inhibition: A Neural Model of Two-Interval Discrimination. Science (80-.). 307, 1121–1124. 10.1126/science.1104171.

8. Fritsche, M., and de Lange, F.P. (2019). Reference repulsion is not a perceptual illusion. Cognition 184, 107–118. 10.1016/j.cognition.2018.12.010.

9. Luu, L., and Stocker, A.A. (2018). Post-decision biases reveal a self-consistency principle in perceptual inference. Elife 7, 1–24. 10.7554/eLife.33334.

10. Zamboni, E., Ledgeway, T., McGraw, P. V, and Schluppeck, D. (2016). Do perceptual biases emerge early or late in visual processing? Decision-biases in motion perception. Proc. R. Soc. B Biol. Sci. 283, 20160263. 10.1098/rspb.2016.0263.

11. Stocker, A.A., and Simoncelli, E.P. (2009). A Bayesian model of conditioned perception. Adv. Neural Inf. Process. Syst. 20 - Proc. 2007 Conf.

12. Luu, L., and Stocker, A.A. (2021). Categorical judgments do not modify sensory representations in working memory. PLoS Comput. Biol. 17, 1–28. 10.1371/journal.pcbi.1008968.

13. Compte, A. (2000). Synaptic Mechanisms and Network Dynamics Underlying Spatial Working Memory in a Cortical Network Model. Cereb. Cortex 10, 910–923. 10.1093/cercor/10.9.910.

14. Schneegans, S., and Bays, P.M. (2018). Drift in neural population activity causes working memory to deteriorate over time. J. Neurosci. 38, 4859–4869. 10.1523/JNEUROSCI.3440-17.2018.

15. Murray, J.D., Jaramillo, J., and Wang, X.J. (2017). Working memory and decision-making in a frontoparietal circuit model. J. Neurosci. 37, 12167–12186. 10.1523/JNEUROSCI.0343-17.2017.

16. Meister, M.L.R., Hennig, J.A., and Huk, A.C. (2013). Signal Multiplexing and Single-Neuron Computations in Lateral Intraparietal Area During Decision-Making. J. Neurosci. 33, 2254–2267. 10.1523/JNEUROSCI.2984-12.2013.

17. Wang, X.J. (2008). Decision Making in Recurrent Neuronal Circuits. Neuron 60, 215–234. 10.1016/j.neuron.2008.09.034.

18. Lemus, L., Hernández, A., Luna, R., Zainos, A., Nácher, V., and Romo, R. (2007). Neural correlates of a postponed decision report. Proc. Natl. Acad. Sci. U. S. A. 104, 17174– 17179. 10.1073/pnas.0707961104.

19. Huttenlocher, J., Hedges, L. V., and Duncan, S. (1991). Categories and particulars: Prototype effects in estimating spatial location. Psychol. Rev. 98, 352–376. 10.1037/0033-295x.98.3.352.

20. Bae, G. (2021). Neural evidence for categorical biases in location and orientation representations in a working memory task. Neuroimage 240, 118366. 10.1016/j.neuroimage.2021.118366.

21. Blake, R., Cepeda, N.J., and Hiris, E. (1997). Memory for Visual Motion. J. Exp. Psychol. Hum. Percept. Perform. 23, 353–369. 10.1037/0096-1523.23.2.353.

22. Rauber, H.J., and Treue, S. (1998). Reference repulsion when judging the direction of visual motion. Perception 27, 393–402. 10.1068/p270393.

23. de Gardelle, V., Kouider, S., and Sackur, J. (2010). An oblique illusion modulated by visibility: Non-monotonic sensory integration in orientation processing. J. Vis. 10, 6–6. 10.1167/10.10.6.

24. Pratte, M.S., Park, Y.E., Rademaker, R.L., and Tong, F. (2017). Accounting for stimulus-specific variation in precision reveals a discrete capacity limit in visual working memory. J. Exp. Psychol. Hum. Percept. Perform. 43, 6–17. 10.1037/xhp0000302.

25. Hardman, K.O., Vergauwe, E., and Ricker, T.J. (2017). Categorical working memory representations are used in delayed estimation of continuous colors. J. Exp. Psychol. Hum. Percept. Perform. 43, 30–54. 10.1037/xhp0000290.

26. Panichello, M.F., DePasquale, B., Pillow, J.W., and Buschman, T.J. (2019). Error-correcting dynamics in visual working memory. Nat. Commun. 10, 1–11. 10.1038/s41467-019-11298-3.

27. Wei, X., and Stocker, A.A. (2015). A Bayesian observer model constrained by efficient coding can explain “anti-Bayesian” percepts. Nat. Neurosci. 18, 1509–1517. 10.1038/nn.4105.

28. Wei, X., and Stocker, A.A. (2017). Lawful relation between perceptual bias and discriminability. Proc. Natl. Acad. Sci. 114, 10244–10249. 10.1073/pnas.1619153114.

29. Hahn, M., and Wei, X. (2024). A unifying theory explains seemingly contradictory biases in perceptual estimation. Nat. Neurosci. 27, 793–804. 10.1038/s41593-024-01574-x.

30. Wimmer, K., Nykamp, D.Q., Constantinidis, C., and Compte, A. (2014). Bump attractor dynamics in prefrontal cortex explains behavioral precision in spatial working memory. Nat. Neurosci. 17, 431–439. 10.1038/nn.3645.

31. Eissa, T.L., and Kilpatrick, Z.P. (2023). Learning efficient representations of environmental priors in working memory. PLoS Comput. Biol. 19, 1–28. 10.1371/journal.pcbi.1011622.

32. Yang, J., Zhang, H., and Lim, S. (2024). Sensory-memory interactions via modular structure explain errors in visual working memory. Elife 13. 10.7554/eLife.95160.4.

33. Stein, H., Barbosa, J., Rosa-Justicia, M., Prades, L., Morató, A., Galan-Gadea, A., Ariño, H., Martinez-Hernandez, E., Castro-Fornieles, J., Dalmau, J., et al. (2020). Reduced serial dependence suggests deficits in synaptic potentiation in anti-NMDAR encephalitis and schizophrenia. Nat. Commun. 11. 10.1038/s41467-020-18033-3.

34. Tomassini, A., Morgan, M.J., and Solomon, J.A. (2010). Orientation uncertainty reduces perceived obliquity. Vision Res. 50, 541–547. 10.1016/j.visres.2009.12.005.

35. Yu, Q., Panichello, M.F., Cai, Y., Postle, B.R., and Buschman, T.J. (2020). Delay-period activity in frontal, parietal, and occipital cortex tracks noise and biases in visual working memory. PLOS Biol. 18, e3000854. 10.1371/journal.pbio.3000854.

36. Girshick, A.R., Landy, M.S., and Simoncelli, E.P. (2011). Cardinal rules: visual orientation perception reflects knowledge of environmental statistics. Nat. Neurosci. 14, 926–932. 10.1038/nn.2831.

37. Taylor, R., and Bays, P.M. (2018). Efficient coding in visual working memory accounts for stimulus-specific variations in recall. J. Neurosci. 38, 7132–7142. 10.1523/JNEUROSCI.1018-18.2018.

38. Ganguli, D., and Simoncelli, E.P. (2014). Efficient Sensory Encoding and Bayesian Inference with Heterogeneous Neural Populations. Neural Comput. 26, 2103–2134. 10.1162/NECO_a_00638.

39. Benjamin, A.S., Zhang, L.Q., Qiu, C., Stocker, A.A., and Kording, K.P. (2022). Efficient neural codes naturally emerge through gradient descent learning. Nat. Commun. 13, 1– 12. 10.1038/s41467-022-35659-7.

40. Mao, J., and Stocker, A.A. (2024). Sensory perception is a holistic inference process. Psychol. Rev. 10.1037/rev0000457.

41. Morais, M.J., and Pillow, J.W. (2018). Power-law efficient neural codes provide general link between perceptual bias and discriminability. Adv. Neural Inf. Process. Syst. 2018-Decem, 5071–5080.

42. Brouwer, G.J., and Heeger, D.J. (2009). Decoding and Reconstructing Color from Responses in Human Visual Cortex. J. Neurosci. 29, 13992–14003. 10.1523/JNEUROSCI.3577-09.2009.

43. Rademaker, R.L., Chunharas, C., and Serences, J.T. (2019). Coexisting representations of sensory and mnemonic information in human visual cortex. Nat. Neurosci. 22, 1336– 1344. 10.1038/s41593-019-0428-x.

44. Harrison, W.J., Bays, P.M., and Rideaux, R. (2023). Neural tuning instantiates prior expectations in the human visual system. Nat. Commun. 14. 10.1038/s41467-023-41027-w.

45. Serences, J.T., Ester, E.F., Vogel, E.K., and Awh, E. (2009). Stimulus-specific delay activity in human primary visual cortex. Psychol. Sci. 20, 207–214. 10.1111/j.1467-9280.2009.02276.x.

46. Master, S.L., Li, S., and Curtis, C.E. (2024). Trying Harder: How Cognitive Effort Sculpts Neural Representations during Working Memory. J. Neurosci. 44, 1–12. 10.1523/JNEUROSCI.0060-24.2024.

47. Kriegeskorte, N., Mur, M., and Bandettini, P. (2008). Representational similarity analysis – connecting the branches of systems neuroscience. Front. Syst. Neurosci. 2, 1–28. 10.3389/neuro.06.004.2008.

48. Friston, K.J., Fletcher, P., Josephs, O., Holmes, A., Rugg, M.D., and Turner, R. (1998). Event-related fMRI: Characterizing differential responses. Neuroimage 7, 30–40. 10.1006/nimg.1997.0306.

49. Yang, G.R., Joglekar, M.R., Song, H.F., Newsome, W.T., and Wang, X.J. (2019). Task representations in neural networks trained to perform many cognitive tasks. Nat. Neurosci. 22, 297–306. 10.1038/s41593-018-0310-2.

50. Dubreuil, A., Valente, A., Beiran, M., Mastrogiuseppe, F., and Ostojic, S. (2022). The role of population structure in computations through neural dynamics. Nat. Neurosci. 25, 783– 794. 10.1038/s41593-022-01088-4.

51. Xie, X., Hahnloser, R.H.R., and Sebastian Seung, H. (2002). Selectively grouping neurons in recurrent networks of lateral inhibition. Neural Comput. 14, 2627–2646. 10.1162/089976602760408008.

52. Birman, D., and Gardner, J.L. (2019). A flexible readout mechanism of human sensory representations. Nat. Commun. 10, 1–13. 10.1038/s41467-019-11448-7.

53. Wolff, M.J., Jochim, J., Akyürek, E.G., Buschman, T.J., and Stokes, M.G. (2020). Drifting codes within a stable coding scheme for working memory. PLoS Biol. 18, e3000625. 10.1371/journal.pbio.3000625.

54. Talluri, B.C., Urai, A.E., Tsetsos, K., Usher, M., and Donner, T.H. (2018). Confirmation Bias through Selective Overweighting of Choice-Consistent Evidence. Curr. Biol. 28, 3128–3135.e8. 10.1016/j.cub.2018.07.052.

55. Glickman, M., Moran, R., and Usher, M. (2022). Evidence integration and decision confidence are modulated by stimulus consistency. Nat. Hum. Behav. 6, 988–999. 10.1038/s41562-022-01318-6.

56. Lange, R.D., Chattoraj, A., Beck, J.M., Yates, J.L., and Haefner, R.M. (2021). A confirmation bias in perceptual decision-making due to hierarchical approximate inference. PLOS Comput. Biol. 17, e1009517. 10.1371/journal.pcbi.1009517.

57. Yang, G.R., Murray, J.D., and Wang, X.J. (2016). A dendritic disinhibitory circuit mechanism for pathway-specific gating. Nat. Commun. 7. 10.1038/ncomms12815.

58. Paninski, L., and Cunningham, J.P. (2018). Neural data science: accelerating the experiment-analysis-theory cycle in large-scale neuroscience. Curr. Opin. Neurobiol. 50, 232–241. 10.1016/j.conb.2018.04.007.

59. van den Berg, R., Shin, H., Chou, W.-C., George, R., and Ma, W.J. (2012). Variability in encoding precision accounts for visual short-term memory limitations. Proc. Natl. Acad. Sci. 109, 8780–8785. 10.1073/pnas.1117465109.

60. Bays, P.M. (2014). Noise in neural populations accounts for errors in working memory. J. Neurosci. 34, 3632–3645. 10.1523/JNEUROSCI.3204-13.2014.

61. Fritsche, M., Mostert, P., and de Lange, F.P. (2017). Opposite Effects of Recent History on Perception and Decision. Curr. Biol. 27, 590–595. 10.1016/j.cub.2017.01.006.

62. Bliss, D.P., Sun, J.J., and D’Esposito, M. (2017). Serial dependence is absent at the time of perception but increases in visual working memory. Sci. Rep. 7, 1–13. 10.1038/s41598-017-15199-7.

63. Markov, Y.A., Tiurina, N.A., and Pascucci, D. (2024). Serial dependence: A matter of memory load. Heliyon 10, e33977. 10.1016/j.heliyon.2024.e33977.

64. Kilpatrick, Z.P. (2018). Synaptic mechanisms of interference in working memory. Sci. Rep. 8, 1–20. 10.1038/s41598-018-25958-9.

65. Barbosa, J., Stein, H., Martinez, R.L., Galan-Gadea, A., Li, S., Dalmau, J., Adam, K.C.S., Valls-Solé, J., Constantinidis, C., and Compte, A. (2020). Interplay between persistent activity and activity-silent dynamics in the prefrontal cortex underlies serial biases in working memory. Nat. Neurosci. 23, 1016–1024. 10.1038/s41593-020-0644-4.

66. Esteban, O., Markiewicz, C.J., Blair, R.W., Moodie, C.A., Isik, A.I., Erramuzpe, A., Kent, J.D., Goncalves, M., DuPre, E., Snyder, M., et al. (2019). fMRIPrep: a robust preprocessing pipeline for functional MRI. Nat. Methods 16, 111–116. 10.1038/s41592-018-0235-4.

67. Gardner, J.L., Merriam, E.P., Schluppeck, D., and Larsson, J. (2018). MGL: Visual psychophysics stimuli and experimental design package (Version 2.0). Zenodo. 10.5281/zenodo.1299497.

68. Gardner, J.L., Merriam, E.P., Schluppeck, D., Besle, J., and Heeger, D.J. (2018). mrTools: Analysis and visualization package for functional magnetic resonance imaging data (Version 4.7). Zenodo. 10.5281/zenodo.1299483.

69. Engel, S.A., Rumelhart, D.E., Wandell, B.A., Lee, A.T., Glover, G.H., Chichilnisky, E.-J., and Shadlen, M.N. (1994). fMRI of human visual cortex. Nature 369, 525–525. 10.1038/369525a0.

70. Choe, K.W., Blake, R., and Lee, S.-H. (2014). Dissociation between Neural Signatures of Stimulus and Choice in Population Activity of Human V1 during Perceptual Decision-Making. J. Neurosci. 34, 2725–2743. 10.1523/JNEUROSCI.1606-13.2014.

71. Ryu, J., and Lee, S.-H. (2017). Stimulus-Tuned Structure of Correlated fMRI Activity in Human Visual Cortex. Cereb. Cortex 28, 693–712. 10.1093/cercor/bhw411.

72. Wang, L., Mruczek, R.E.B., Arcaro, M.J., and Kastner, S. (2015). Probabilistic maps of visual topography in human cortex. Cereb. Cortex 25, 3911–3931. 10.1093/cercor/bhu277.

73. Glasser, M.F., Coalson, T.S., Robinson, E.C., Hacker, C.D., Harwell, J., Yacoub, E., Ugurbil, K., Andersson, J., Beckmann, C.F., Jenkinson, M., et al. (2016). A multi-modal parcellation of human cerebral cortex. Nature 536, 171–178. 10.1038/nature18933.

74. Lindquist, M.A., Geuter, S., Wager, T.D., and Caffo, B.S. (2019). Modular preprocessing pipelines can reintroduce artifacts into fMRI data. Hum. Brain Mapp. 40, 2358–2376. 10.1002/hbm.24528.

75. Rademaker, R.L., Bloem, I.M., De Weerd, P., and Sack, A.T. (2015). The impact of interference on short-term memory for visual orientation. J. Exp. Psychol. Hum. Percept. Perform. 41, 1650–1665. 10.1037/xhp0000110.

76. Green, D.M., and Swets, J.A. (1966). Signal Detection Theory and Psychophysics (Wiley).

77. Glover, G.H. (1999). Deconvolution of Impulse Response in Event-Related BOLD fMRI. Neuroimage 9, 416–429. 10.1006/nimg.1998.0419.

78. Walther, A., Nili, H., Ejaz, N., Alink, A., Kriegeskorte, N., and Diedrichsen, J. (2016). Reliability of dissimilarity measures for multi-voxel pattern analysis. Neuroimage 137, 188–200. 10.1016/j.neuroimage.2015.12.012.

79. Pietrini, P., Furey, M.L., Ricciardi, E., Gobbini, M.I., Wu, W.H.C., Cohen, L., Guazzelli, M., and Haxby, J. V. (2004). Beyond sensory images: Object-based representation in the human ventral pathway. Proc. Natl. Acad. Sci. U. S. A. 101, 5658–5663. 10.1073/pnas.0400707101.

80. Risken, H. (1996). The Fokker-Planck Equation: Methods of Solution and Applications (Springer).

## Supplemental References

[S1] Zamboni, E., Ledgeway, T., McGraw, P.V., and Schluppeck, D. (2016). Do perceptual biases emerge early or late in visual processing? Decision-biases in motion perception. Proc. R. Soc. B Biol. Sci. 283, 20160263. doi: 10.1098/rspb.2016.0263.

[S2] Fritsche, M., and de Lange, F.P. (2019). Reference repulsion is not a perceptual illusion. Cognition 184, 107–118. doi: 10.1016/j.cognition.2018.12.010.

[S3] Rademaker, R.L., Bloem, I.M., De Weerd, P., and Sack, A.T. (2015). The impact of interference on short-term memory for visual orientation. J. Exp. Psychol. Hum. Percept. Perform. 41, 1650. doi: 10.1037/xhp0000110.

[S4] Rademaker, R.L., Chunharas, C., and Serences, J.T. (2019). Coexisting representations of sensory and mnemonic information in human visual cortex. Nat. Neurosci. 22, 1336–1344. doi: 10.1038/s41593-019-0428-x.

[S5] Ganguli, D., and Simoncelli, E.P. (2014). Efficient sensory encoding and bayesian inference with heterogeneous neural populations. Neural Comput. 26, 2103–2134. doi: 10.1162/NECO_a_00638.

[S6] Wei, X.X., and Stocker, A.A. (2015). A Bayesian observer model constrained by efficient coding can explain ‘anti-Bayesian’ percepts. Nat. Neurosci. 18, 1509–1517. doi: 10.1038/nn.4105.

[S7] Taylor, R., and Bays, P.M. (2018). Efficient coding in visual working memory accounts for stimulus-specific variations in recall. J. Neurosci. 38, 7132–7142. doi: 10.1523/JNEUROSCI.1018-18.2018.

[S8] Liu, T., Cable, D., and Gardner, J.L. (2018). Inverted encoding models of human population response conflate noise and neural tuning width. J. Neurosci. 38, 398–408. doi: 10.1523/JNEUROSCI.2453-17.2017.

[S9] van Bergen, R.S., Ma, W.J., Pratte, M.S., and Jehee, J.F. (2015). Sensory uncertainty decoded from visual cortex predicts behavior. Nat. Neurosci. 18, 1728–1730. doi: 10.1038/nn.4150.

[S10] Zhang, R.Y., Wei, X.X., and Kay, K. (2020). Understanding multivariate brain activity: Evaluating the effect of voxelwise noise correlations on population codes in functional magnetic resonance imaging. PLoS Comput. Biol. 16, e1008153. doi: 10.1371/journal.pcbi.1008153.

[S11] Girshick, A.R., Landy, M.S., and Simoncelli, E.P. (2011). Cardinal rules: visual orientation perception reflects knowledge of environmental statistics. Nat. Neurosci. 14, 926–932. doi: 10.1038/nn.2831.

[S12] Yang, J., Zhang, H., and Lim, S. (2024). Sensory-memory interactions via modular structure explain errors in visual working memory. Elife 13, RP95160. doi: 10.7554/eLife.95160.3.

[S13] Hahn, M., and Wei, X.X. (2024). A unifying theory explains seemingly contradictory biases in perceptual estimation. Nat. Neurosci. 27, 793–804. doi: 10.1038/s41593-024-01574-x.

